# Uncovering the Basis of Human Connectome Complexity: The Role of Neuronal Morphology

**DOI:** 10.64898/2026.02.12.705493

**Authors:** Natalí Barros-Zulaica, Daniela Egas Santander, Lida Kanari, Ying Shi, Rodrigo Perin, Maurizio Pezzoli, Ruth Benavides-Piccione, Javier DeFelipe, Christiaan P J de Kock, Idan Segev, Henry Markram, Michael W. Reimann

## Abstract

Comparative studies have established differences between the electrophysiology and anatomy of human and rodent cortical circuits. A consistent finding is that human neuronal morphologies display more elaborate neurite shapes than those of rodents, a feature that cannot be accounted for merely by their larger size according to recent findings. Here, we study the impact of these neurite shapes on the structure of synaptic connectivity in their local microcircuitry. Our approach is based on the idea that axonal and dendritic geometries constrain the locations of afferent and efferent synaptic contacts (potential connectivity). Although the mechanisms by which potential connectivity translates into actual synaptic connectivity are manifold and complex, the potential connectivity is nevertheless highly informative for the final structure of a biological connectome. We found that connectomes predicted from human reconstructed morphologies have higher complexity according to several measures that have been demonstrated to be functionally relevant. Going beyond a simple comparison, we demonstrate mechanistically how the shapes of neuron morphologies give rise to non-random and clustered structures observed in experimentally measured connectomes, and how the specific shapes of human neurons strengthen the process. Finally, we conceptually examine how synapse formation processes may interact with potential connectivity, showing that a process compatible with Hebbian plasticity leads to the highest complexity and best match experimentally observed patterns.

## 1 Introduction

It is believed that the neocortex is the brain structure that most differentiates humans from other mammals, as its activity is related to the cognitive skills thought to be uniquely human, such as spoken language, writing, composing music, or inventing complex machines [1, 2].

Starting with Santiago Ramon y Cajal’s anatomical and histological inquiries in the 19th century [3], many studies have been performed to unravel the anatomical structure of the neocortex of several species [1, 4]. Comparative studies have described structural and functional rules that are common between different mammal species, but also found specific characteristic differences between them, e.g., between humans and rodents [1, 4–6]. For instance, it has been shown that the human cortex has lower cell density [1, 7], higher ratios of inhibitory cells over excitatory cells [7, 8], as well as stronger and more reliable synapses [5, 9–11].

With respect to the anatomy of cortical connectivity, several studies have measured and quantified the local connectivity in the human cortex [12, 13]; as well as compared the connectivity properties of humans and mice [7, 14]. However, open questions about the sources of differences in connectivity between species remain open.

Questions that are difficult to answer with purely experimental methods have been successfully addressed through the use of computational models. Depending on the question, this can require simplified [15, 16], or morphologically and biophysically detailed models [17–20]. Models generate specific predictions that can be tested in the future with experimental techniques. For example, to predict the number of vesicles ready to be released at the presynaptic site of different neuron types in the rat somatosensory cortex [21], or understanding the diversity of neural code through network connectivity and activity [22].

Neuron morphology impacts network structure at different computational scales [23–26], and we hypothesize that it also does so at the micro-scale, and in an explainable manner. We argue that differences in morphology between species can be used to predict corresponding differences in their networks and their theoretical computational capabilities. In this regard, Cragg [27] hypothesized: “*works of Nissl [28] on the greater separation of human neurons, compared with those of mole or dog brain, and the remarks of von Economo [29] on the greater opportunity for neuronal interaction provided by the more extensive fibre plexus that fills the space between the widely separated neurons*.”

Furthermore, in a comparative study between species [30], differences in neuronal shape and dendritic complexity between homologous regions of the human and mouse brain were identified and linked to the topology of the corresponding neuronal networks. Human cortical pyramidal cells exhibit more complex dendritic arborization, in terms of topological properties [31], resulting in higher computational capacity and more complex topological structures in human networks.

However, Kanari et al. [30] considered the potential connectivity that arises from locations where axons and dendrites meet, randomly reducing the number of touches until the biological number of synaptic contacts present in actual cortical circuits was reached [32]. In this paper, we extend that work in three ways. First by using a broader network analysis methods, which allowed us to compare and validate the networks’ structure to experimental results at different scales [12, 13]. Second by providing a mechanistic explanation of the source of the differences originating from the neuronal morphologies. Third, by exploring a variety of methods to select a subset of potential synaptic contacts other than selecting a random subset. This allowed us to investigate to what degree the selection of active synapses can be described by simple stochastic processes, and how much specificity for cell types or subcellular domains is required [33–35], which remains a point of contention in the literature [36].

We used the extensively validated models of rat somatosensory cortex of Reimann et al. [34], Isbister et al. [37] and a recent, biologically detailed model of human temporal cortex built according to similar methods [11]. We focused our study on the connectivity between layers 2 and 3 pyramidal cell networks for which the human data is more abundant [9, 14, 38, 39]. To enable meaningful comparison, we extracted central subvolumes of approximately 20000 neurons for each of the models to perform connectivity analyses.

Our main findings are as follows. First, the method used for selecting a subset of potential synapses drastically affects the structure of connectivity and difference between human and rat models. Although we ensured that the competing methods resulted in the same densities of synapses (touches) or connections (edges), when simulating multi-patch experiments the estimated connection probabilities varied strongly between methods. In particular, we found that an algorithm based on cooperative synapse formation [17, 34] most accurately reproduced connectivity properties observed in experimental setups [12, 13]. Crucially, it also increased the difference between human and rat model with respect to measurements of network complexity. Second, network metrics such as connection probabilities or synapse densities, are strongly affected by the spatial scale of the network (its volume in space and the number of neurons). The effect of spatial scale was stronger for the human model. This also cautions against extrapolating estimates from multi-patch experiments to larger networks. Third, with respect to the substantial differences in connectivity derived from human vs. rodent cell density and morphology, it is specifically the shape of morphologies that matters more than high-level differences, such as lower cell density and larger sizes of human morphologies. Fourth, we provide a more mechanistic explanation of how the complexity of dendritic/axonal trees give rise to complex, structured networks. Taken together, the greater branching complexity of human neurons, compared to those of rodents, gives rise to increased topological complexity in their corresponding networks.

## 2 Results

### 2.1 Morphology differences lead to higher network complexity in human compared to rodent

As mentioned in the introduction, our departure point is the study of Kanari et al., 2025 [30], who observed that a network built from individual mouse morphologies scaled to the human volume, did not result on the same network connectivity complexity as the one from human. In this study we dived deeper and try to explain the reasons for this surprising difference. While in Kanari’s study the connectivity of the networks reflected potential appositions [30], i.e., the number of appositions between axons and dendrites was randomly reduced until a biological connectivity density was achieved, in this study we explored more complex prediction methods and analyze how they affect the difference between rat and human. Additionally, we extended the concept of scaling elements from one species to another by scaling the statistics of the connectivity of the rat model to the human volume.

Specifically, we compared and contrasted three different predicted networks. The first was built from 366 unique reconstructions of Layer 2/3 Pyramidal Cells from human temporal cortex [11] with connectivity based on axo-dendritic touches, filtered by different algorithms that will be described later. The second was a sub-volume representing layers 2/3 of the model of rat somatosensory regions of [17], with connectivity predicted according to the same algorithms as for the human model. The third was a statistical model describing what would happen if human neurons were simply rat neurons scaled in all three dimensions, which we denoted as the *human model based on rat data*(HC_RD). In this model the connection probability of the rat network, considering all potential connections, was scaled according to the differences in cell density, neurite reach and neurite lengths (see methods). We could contrast the first and the third model to understand the effect of specific human morphology shapes on connectivity, since the latter does not take them into account.

We began by confirming that with these models we can reproduce the observations of Kanari et al. [30]. To that end, we used the same algorithm to detect and then filter the axo-dendritic touches as in that study. Touches were detected as axon-dendritic appositions with a maximum distance of 2*µm*. For randomly chosen pairs of neurons, all touches from one to the other were discarded, and the remaining touches were considered as predicted synaptic connections. 96% of connections were removed this way, until the average degree of the network was compatible with biological observations. We called this algorithm “edge to edge pruning”, and the resulting network “E_to_E”, as edges were removed from the network formed by all touches to match a target number of edges. This was applied to both human and rat models (Fig. 1A).

**Figure 1.**
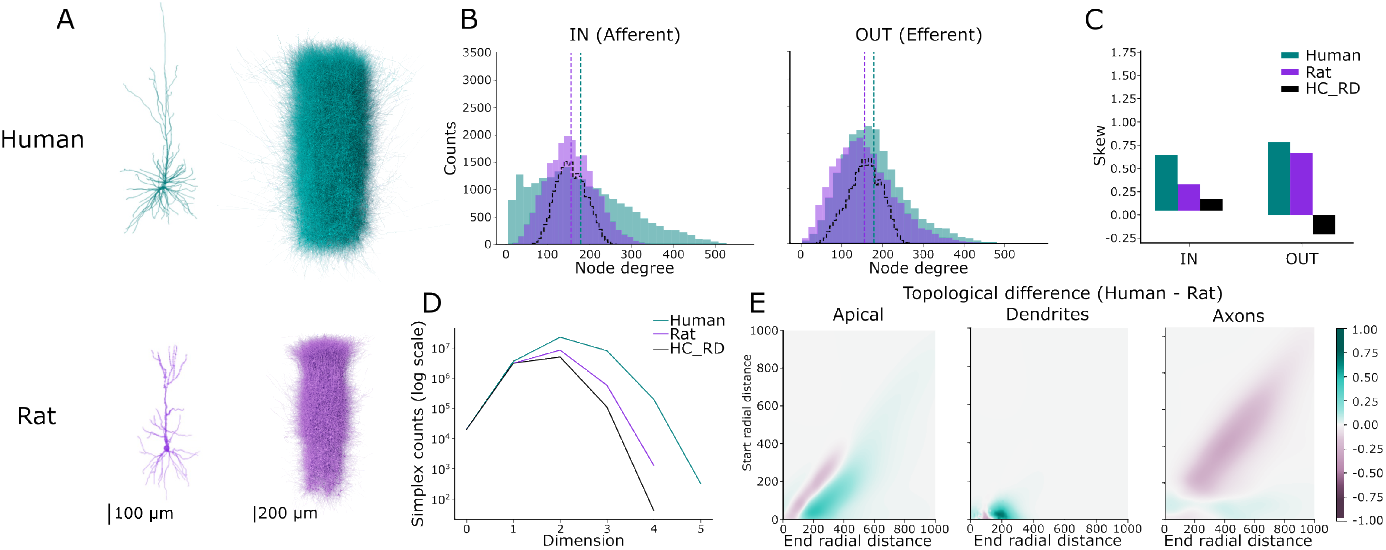
Scaling rat connectivity statistics to human volume showed the same results as Kanari et al., 2025. (A, left) Morphological reconstructions of two cortical pyramidal cells of layer 2/3 in Human (top, teal) and rat (bottom, purple). (A, right) Columns of the cortical microcircuits models of human (top, teal) and rat (bottom, purple). (B) Degree distributions of IN (afferent) (left) and OUT (efferent) (right) connections per node for human (teal), rat (purple) and HC RD model (dashed black line). Dashed vertical lines denote mean values of the distributions accordingly. (C) Skew analysis for the degree distributions in B. (D) Simplex counts in logarithmic scale for human (teal), rat (purple) and HC RD (black). (E) Difference between the respective persistent images for the different neurites: apical dendrites (left), basal dendrites (middle) and axons (right). X-axis represent the start radial distance of branches, Y-axis the end radial distance of branches. Green represents over-expression of human and purple over-expression of rat branches

To ensure comparable results while addressing these questions, we considered circuits on similar numbers of neurons (20072 for human; 20064 for rat), which is different than selecting networks on a similar spatial volume due to the different cell densities. We considered all connectomes as *networks*, with *nodes* (neurons) and the set of *touches* from one node to another forming a (weighted, directed) *edge* (see Fig. 2A; nodes: black dots; touches: red dots; edges: blue lines).

**Figure 2.**
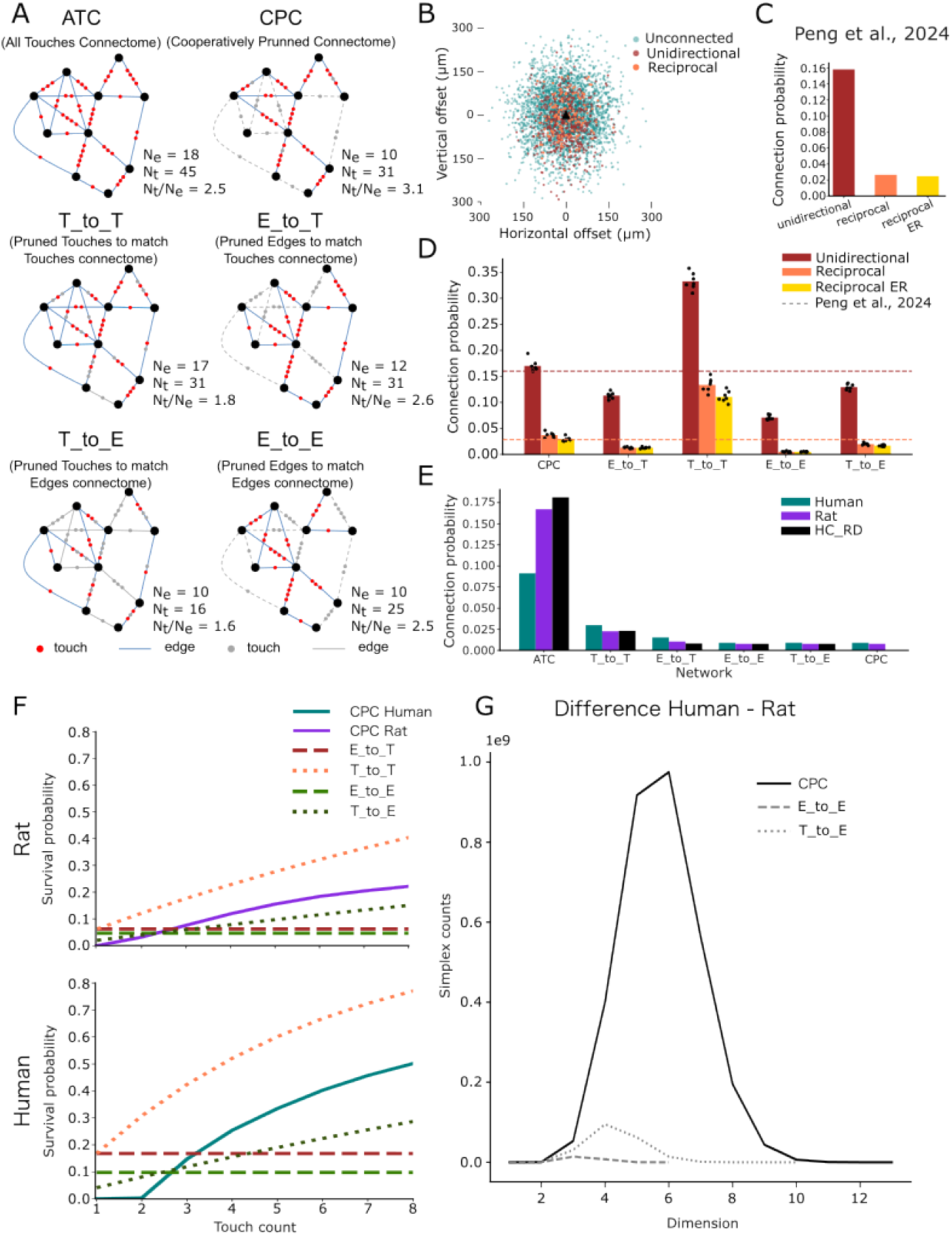
Testing different pruning methods showed differences in network complexity. (A) Diagram showing a toy example of a network with all nodes (black dots) connected (edges, blue lines) through one ore several touches (red dots) (ATC) and its result connectivity after applying different pruning methods: predicted cooperative connectivity using Reimann et al. [32] (CPC); pruning touches(or edges) randomly of the ATC to match the number of touches(or edges) (T(E) to T(E)). The elements of the diagrams are: neurons (nodes, black dots), connections between neurons (edges, blue lines), actual contact points between neurons (touches, red dots), pruned touch (gray dots) and pruned edge (gray lines). *N*_*e*_: total number of edges; *N*_*t*_: total number of touches. (B) Exemplar scatter plot showing the probed (light teal dots), connected (brown dots) and reciprocal (coral dots) connections at certain inter-somatic distances during an *in silico* patch-clamp experiment performed in the human CPC. Pre-synaptic cell soma (black triangle) aligned to origin. (C) Bar plot from Peng et al. [12] Figure 1B. (D) Bar plot showing the connection probability of unidirectional (brown), reciprocal (coral) and reciprocal ER (gold) connections for the human circuit with different pruning methods. Dashed lines point to the values in bar plot B for unidirectional (brown) and reciprocal (coral) connections. (E) Connection probability comparison of rat (purple), human (teal) and the analytical model HC RD (black) for the ATC and for all the different pruning methods.(F) Edge survival probability for the CPCs (solid lines) and the other pruned networks (dashed lines) for rat (top) and human (bottom). (G) Distribution of simplex dimentions for the CPCs (solid black line) and the two pruned networks with the closest connection probabilities to the CPC network: E_to_E (dashed, gray) and T_to_E (dotted, gray).

Using this approach we observed in the E_to_E networks asymmetric distributions for both in- and out-degrees with a heavier right tail, but not for the HC RD control (Fig.1B). The human network showed a higher heterogeneity than the rat, especially for in-degree distributions, resulting in higher skewness around 0.75 (Fig.1C). We concluded that node degree distributions of human networks are more heterogeneous than those of rat, in line with Kanari et al. [30], and that the heterogeneity of the modeled networks couldn’t be matched by the HC_RD control.

To further confirm the differences in network complexity across species of Kanari et al., 2025 [30], we counted larger motifs, specifically directed simplices (Fig.1D). These are structures that represent motifs composed of an increasing number of nodes. A directed n-simplex is a motif on n+1 nodes all-to-all connected in feed-forward fashion; we refer to n as its dimension. In particular, a 0-simplex is a node, a 1-simplex is a directed edge and a 2-simplex is what is sometimes referred to as a transitive triad (Fig.S2F). The counting of simplex over-expression, and in particular determining the maximum dimension attained, has been shown to serve as a metric of network complexity [40–42]. Moreover, simplex over-expression has been observed across species and scales [43–47] and these motifs have been shown to have functional implications [22, 48, 49]. Consequently, we counted the numbers of simplices in each dimension for the human, rat and HC RD networks, finding simplices up to dimension 4 in the rat and HC RD, whereas the human reaches dimension 5 (Fig.1D).

Then, to characterize and compare the branching properties of neurons across species, we used a topological analysis, based on the Topological Morphology Descriptor (TMD) [31]. Using the TMD algorithm one can encode the topology of the branching structure of a neuron as an image (Fig.S5, bottom left). Intuitively, in this image, each pixel’s x-axis represents the distance to the soma and its distance from the diagonal, the length of a branch. Each pixel’s color intensity is determined by the density of branches at that location (see Method for a precise definition). This results in an image summary of the topological complexity of the branching pattern. This image-based topological summary [50] has been shown to be a suitable format for quantitative analysis and has been used in several applications, including neuronal morphology classification [31, 51] and glia classification [52].

We extracted the TMD of human and rat dendrites and quantified their differences. The analysis revealed a higher density of dendritic branches near the soma in human neurons (Fig.1E), consistent with previous findings comparing human and mouse morphologies [30]. For axons, the TMD analysis indicated that rat axons generally extend further (Fig.1E, right), which likely reflects the higher reconstruction quality of rat axonal morphologies relative to the limitations inherent in human axon reconstructions. Nevertheless, we observed an increased axonal branching density near the soma in human neurons, mirroring the dendritic pattern and suggesting a species-specific specialization in proximal arborization.

In summary, the qualitative differences found in Kanari et al. [30] between human and mouse were reproduced between human and rat. More precisely, we observed that the human network complexity could not be reached by the rat network nor by the HC_RD model.

### 2.2 Higher specificity and robustness of human connections enhance network complexity

In the previous section, we confirmed the result that networks predicted from human morphologies result in more complex networks than networks predicted from rodent morphologies. We found this to be true already for a simple algorithm for predicted connectivity inspired by “Peters’ rule”, and for a range of measures of complexity. A control that scaled the distance-dependent statistics of rat connectivity to human network size did not capture the increase in complexity. Hence, it must be related to the specific shape of human neuron morphologies. To better understand which characteristics of human neuron morphologies are responsible, we stepped through the process of connectome prediction of the models ([34] and [11]) in detail and explored alternative algorithms.

Connectivity was predicted by detecting axo-dendritic touches, then filtering the set of touches (we refer to this step as the “pruning step”), and considering the remaining ones as predicted synaptic contacts. The way of pruning may affect the network’s structure and studying the differences may help us to unravel the effect of morphologies on the network topology. We explored six widely used pruning methods [53] (Table S1), resulting in different networks that we could then compare. For the first network, we considered all axo-dendritic touches as predicted synapses, i.e., used no filtering at all, generating what we denoted as the *All-touches connectome (ATC)* (Fig. 2A, top left). Second, we used the algorithm of Reimann et al. [32], parameterized as in Reimann et al. [34], generating what we called the *Cooperatively Pruned Connectome (CPC)* (Fig.2A, top right), as it employs the principle of cooperative synapse formation [54]. Four further pruning methods we used CPC as a reference. The first of them was edge to edge pruning, already employed in (Fig. 1). That is, edges of the network graph were randomly removed to match the number of edges of the graph of the CPC network. The other three used the fact that edges of the network graphs could represent multiple touches from one axon onto the dendrites of another neuron (Fig. 2A, red dots). In “touch to touch” pruning (T to T), individual touches were randomly removed until the number of touches of the CPC was matched. An edge was removed only when its last touch was pruned. “touch to edge” pruning (T to E) thusly removed touches until the number of edges of the CPC was matched; finally “edge to touch” pruning removed edges to match the number of touches.

We began by evaluating the different pruning methods against the experimentally sampled human local connectivity of Peng et al. [12]. We replicated their *in vitro* experiment *in silico*, by sampling pairs of neurons within a maximum range of 300*µm*, but mostly within 150*µm* in all networks (see Methods; Fig.2B, C; compare to Fig.2A in Peng et al. [12]). In agreement with the experimental study, we found, for all pruning methods, the absence of an over-expression of reciprocal connectivity (Fig.2D). However, the connection probabilities differed drastically between them, and CPC was the model that most closely resembled the *in vitro* results. Furthermore, the CPC was the only pruning method that matched the mean number of touches per connection of the biological references for synapses per connection: 4.0 ± 1.2 for human [9], 2.8 ± 0.7 for rat [55] (Fig.S1, top). This supported the use of the CPC as a fair approximation of the experimentally observed connectome and thus, our continued use of it as a reference in this study.

We compared the overall density of edges in the ATCs, i.e., not by sampling pairs, but based on all pairs in the models. We found only around half the density for human compared to rat and an even higher density for HC RD (human: 0.091; rat: 0.17; HC RD: 0.18). Surprisingly, we found for all pruned networks a sudden inversion of connection probabilities, where now the human model is slightly higher, even above HC RD (Fig.2E). This is due to fewer touches being pruned in human than in rat or the HC RD ATC networks across pruning methods (Fig.S1, middle and bottom, Table S2). Note that the amount pruned is not arbitrary, but directly determined by experimental measurements of axonal bouton densities for CPC, and thus also for all other pruning methods. These values are comparable in human and rat, but in human the neuron morphologies and the distances between neurons are larger [11]. This means that the same bouton density represents a higher number and a higher density of connections. This shows that human connections are more robust i.e., they have higher probability of survival across pruning methods. Note in particular, that the connection probability was similar to the rat when comparing the CPCs (human: 0.0089; rat: 0.0078) and that the T to T and E to T networks had higher connection probabilities for all models.

It may appear surprising that different stochastic pruning algorithms can lead to such large systematic differences between the resulting networks. We explain this as follows: While the pruning algorithms are stochastic, they implement different *survival probability functions*, i.e., the survival of a connection in this process depends on its strength, i.e., its number of touches in the ATC, and the shape of the dependence differs between algorithms. Indeed, when evaluated, we found these functions to have very different shapes (Fig.2F). By construction, they were flat for the pruning methods that removed edges randomly while they were increasing for the ones that removed touches randomly and the CPC. Moreover, the CPC curves for both species were steeper than their corresponding T to E curves, and in the case of human more than two touches were needed in order to keep an edge after pruning, but for touch counts above two the human curve was steeper than the curve for rat (Fig.2F). It is important to note that it is this specific shape of the survival function that allows the CPC to simultaneously match biological reference data on edge density (Fig.2D), touches per connection (Fig.S1), and - by construction - synapses per neuron [32]. Conversely, T to E pruning had the same edge density by construction, but admitted more connections formed by one or two touches and fewer formed by more than three (Fig.2F), resulting in fewer touches per connection than the reference (Fig.S1). Hence, the touch count-dependent survival function is not just an observation about an imperfect connectome pruning algorithm, but a biological prediction, based on experimental data. From this prediction we observed: First, that the survival function for CPC in human leads to a network with a more robust and specific set of edges than alternative pruning methods. Second, that this difference is less pronounced in the rat model, where CPC is more of an intermediate between T_to_E and T_to_T.

To assess the consequence of these differences with respect to network complexity, we once again referred to the measure of simplex counts (Fig.2G). We have already demonstrated that for the human model there was higher count of directed simplices of dimensions two and above compared to the rat model, despite both models having approximately the same number of nodes (neurons) and edges (Fig.1D). When contrasting this measure between pruning algorithms, we found that CPC drastically increased this difference (Fig.2G).

Taken together, we found that the results of Kanari et al. [30] are only the starting point for the exploration of how neuronal anatomy leads to differences in network topology between rodent and human. Specifically, we found that the edge-pruning method must be considered to understand the differences. While even the CPC network is certainly imperfect, it simultaneously matches many measures of a reference network and hence can be used for statistical predictions. In that regard, we predicted: First, that the increased complexity of human compared to rat networks is larger than what was found for human versus mouse by Kanari et al. [30]. Second, that this is partially a consequence of higher specificity and robustness for human connections, i.e., the survival function (Fig.2F) increased more sharply and reached higher values. In other words, the human cortex could sustain a larger number of synapses, relative to the overall anatomical scale, compared to the rat. Third, that in this process individual neuron shape matters, as the HC_RD control was closer to rat than to human (Fig.2E).

### 2.3 Human neuronal networks are more strongly characterized by distance-dependence than rat networks

One complication of comparing human to rat local circuitry is their difference in spatial scale. To ensure comparable graph-level statistics, we chose to compare networks with approximately the same number of neurons. However, this resulted in models of different spatial sizes (Fig.1A). This was a potentially confounding factor, as connection probability is widely known to be distance-dependent [56]. For this reason, we explored and compared the distance-dependence of our model networks.

For the ATC networks we found almost identical profiles of distance-dependent connection probability between human and rat (Fig.3A, top, teal vs. purple). This was a surprise, as the overall larger spatial scale of human neurons would imply a higher spatial range. Indeed, the HC_RD network had a less steep decrease with distance (Fig.3A, top, black). Furthermore, for CPC the human network had a steeper decrease than the rat network (Fig.3A, bottom). This means that human networks will be more strongly characterized and shaped by distance-dependence.

**Figure 3.**
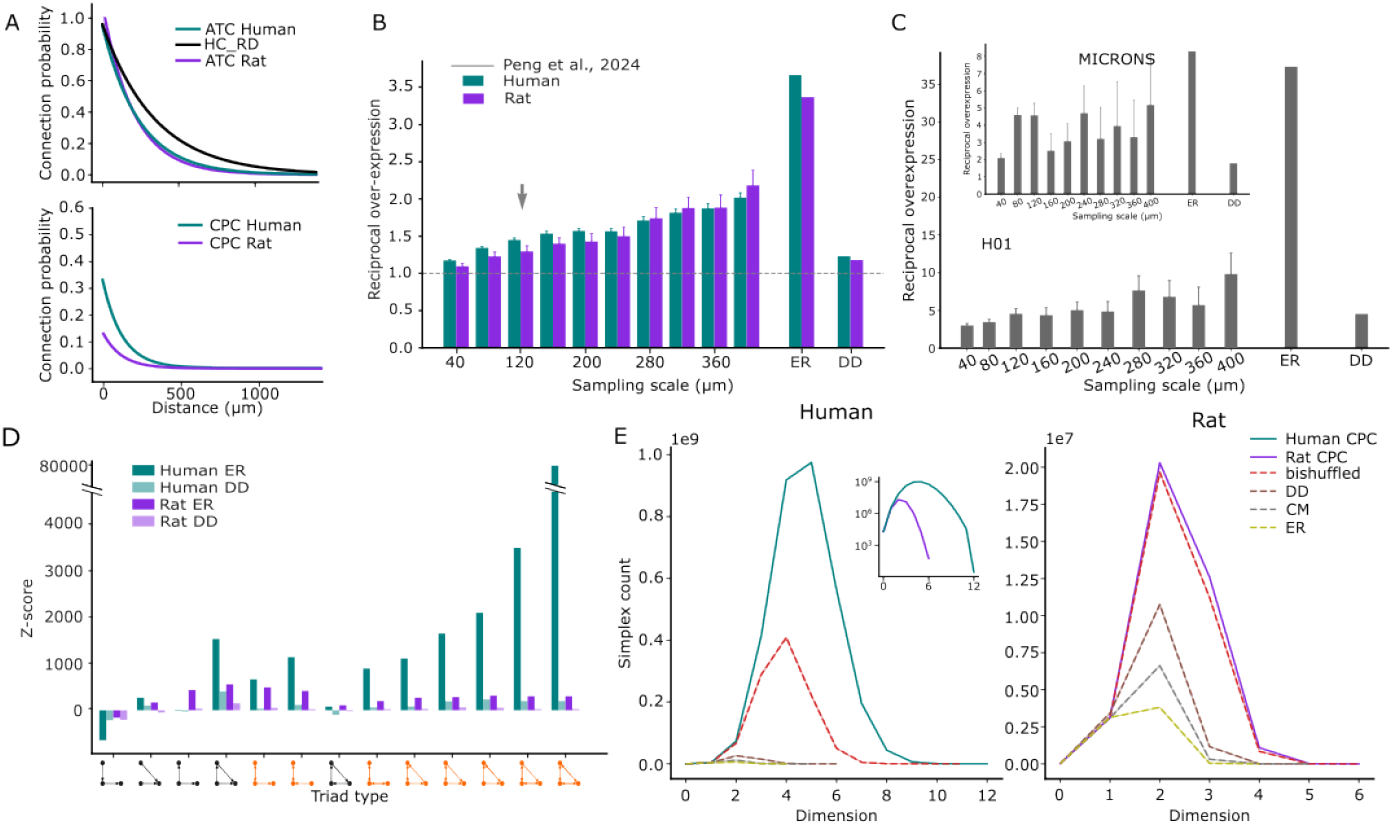
The size of the sampling scale considered affects network complexity measurements. (A) Connection probability along distance for the ATCs (top) and the CPCs (bottom) of human (teal), rat (purple) and the HC_RD (black). (B) Bar plot representing the over-expression of reciprocal connections at different sampling scales from (40 - 400) µm for the CPCs and two random models of Erdös-Rényi (ER) and distance-dependent (DD), for human (teal) and rat (purple). Dashed gray line and gray arrow represents the value of reciprocal over-expression and the sampling scale for Peng et al., [12] respectively. (C) Over-expression of reciprocal connections at different scales (40 and 400 µm) for the H01 [13] (bottom) and MICRONS [57] (top) datasets. (D) Normalized representation of triad motifs for full scale CPC networks for human (teal) and rat (purple). Normalized against two random models ER (dark colors) and DD (light colors). Triad motives in orange contain reciprocal connections. (E) Distribution of simplex dimension for the CPCs (solid lines) and some controls (dashed lines): bishuffled (red), T to E (violet), distance dependent (DD; brown), E_to_E (pink), configuration model (CM; gray) and Erdös-Rényi (ER; green), for human (left) and rat (right). Subplot: same as main plot for logarithmic scale of human and rat CPCs.

To understand the consequences of this, we investigated how the distance-dependence affected the non-random structure of human and rat network models. To that end, we repeated the *in silico* patch-clamp experiment we used to validate the human CPC against the data of Peng et al. [12] (Fig.2B, C), but at increased or decreased spatial scales. Previously, the horizontal and vertical offsets of sampled neuron pairs followed a 2d Gaussian distribution with a standard deviation of 120*µm* (Fig.2B). We varied that standard deviation between 40 and 400*µm*, thereby focusing on pairs at different distances. We called that standard deviation the “spatial scale” of the sampling experiment. For a scale of 120*µm* that best matched the sampling of Peng et al. [12] we approximately matched their reported connection probability and reciprocal connection probability (Fig.2C). Consequently, we approximately matched their finding that reciprocal connectivity was not (or in our case: only modestly) overexpressed compared to an Erdős–Rényi control (Fig.3B, grey arrow). However, we found that this measure strongly increased with spatial scale, reaching values around 2.0 at 400*µm*. This demonstrates that spatial scale does indeed affect measures of non-random structure. This was further confirmed when we calculated reciprocal over-expression for all pairs, i.e., without patch-sampling (around 20, 000^2^ pairs): While relative to an Erdős–Rényi control we found a 3.5-fold over-expression, this was reduced to a modest 1.2 relative to a control that captured the distance-dependence of connection probability (Fig.3B, right).

With respect to human versus rat, we found no significant differences in the CPC networks for this analysis (Fig.3B, teal vs. purple). However, this differed when we instead performed this analysis on connectomes reconstructed with electron microscopy, specifically the connectome of adult mouse V1 of (*MICrONS* [57]) and the connectome of adult human temporal cortex of (*H01* [13]). For these datasets, we found a more profound increase of reciprocal overexpression with sampling scale for human (Fig.3C). Additionally, for MICrONS the reciprocal over-expression relative to Erdős–Rényi was four times higher than the one relative to a distance-dependent control, whereas for H01 this value was seven times higher. This confirms our prediction that human networks are more strongly characterized and shaped by distance-dependence.

We extended this type of analysis further, moving from reciprocal connectivity to an analysis of triad motif over-expression. We counted triad motifs in the entire CPC networks for human and rat and normalized the resulting counts against Erdős–Rényi and distance-dependent controls by subtracting the expected mean counts and dividing by the expected standard deviation, i.e., we calculated a z-score (Fig.3D). We found strong over-expression in both models and for both normalization methods. We found over-expression for all motifs with more than two connections, except for the cycle motif. For over-expressed motifs, the level of over-expression was consistently higher for the human compared to the rat model. Crucially, the difference between the normalization methods, i.e., against Erdős–Rényi versus a distance-dependent model, was larger for human than for rat. This once again indicates that the structure of the human CPC network is more strongly affected by distance-dependence. However even when normalized against the distance-dependent control, over-expression was still stronger for the human model.

Curiously, the highest levels of over-expression in human were found for motifs that included reciprocal connections (Fig.3D, orange triads). This indicates that reciprocal connections in the human CPC network are found specifically within strongly inter-connected motifs. That is, while reciprocal connections are globally only mildly over-expressed (and not more strongly than in rat CPC), their locations within the topology of the network are strongly non-random. To further confirm this trend, we defined a new control network, the *bishuffled* control. It keeps the undirected structure of the original network identical, but randomizes the location of reciprocal connections. If the value of a network measure differs between original network and this control, then we can conclude that the location of reciprocal connections is non-random with respect to that measure. Indeed, we found for directed simplex counts in the human CPC network a large difference to the bishuffled control. It was much larger than for the rat CPC network (Fig.3E). This indicates that the probability that a connection between two neurons is reciprocal is mediated by the participation of that connection in simplicial motifs.

Taken together, we predicted that human neuronal networks are characterized more strongly by distance-dependence than rat networks. First, over the range of inter-soma distances that are observed in local circuitry, there is a sharper decrease in connection probability for human than for rat. Second, this means that the spatial scale at which neuronal pairs are sampled when connectivity is measured can affect the results. The results of such experiments are often analyzed under the assumption that distance-dependence has no effect at the sampled scales, e.g., comparing results to Erdős–Rényi controls. We caution against this, in particular for human circuitry. As a corollary, this may explain the result of Peng et al. [12] that human circuitry does not over-express reciprocal connections. We suggested that the outcome may differ had pairs been sampled at larger distances. Third, even if reciprocal connections are not more abundant than expected, their locations within the network are non-random. They are structurally located in a way that substantially increases simplex motif counts. These points once again demonstrate ways in which human neuronal morphologies lead to more complex network structure.

### 2.4 Spatial neurite variability provides the backbone for the differences between human and rat

Thus far, we have demonstrated several ways in which networks formed from human neuronal morphologies are more complex than networks formed from rat morphologies, and that this is related to the shape of individual morphologies. In this last section we explore which specific features of the morphologies explain these results.

We have previously shown (Fig.2F) that for human networks a different connection survival probability function was observed. Input into this function was the number of touches that form a potential connection. We therefore examined the statistics of how the touches, and especially multiple touches for a pair, were formed. One hypothesis was that touches in different parts of the dendritic and axonal tree are formed independently, giving rise to a geometric or binomial distribution of touch counts. However, we found that this process was not statistically independent at all. Instead, we found that the presence of an axo-dendritic touch drastically increased the probability of finding additional ones, and this increase grew stronger and stronger with the number of confirmed touches (Fig.4A). Distance-dependence cannot explain this, as it was observed across different distance bins. Crucially, for rat ATC this effect only appeared for neuron pairs further than 100*µm* apart, whereas for human ATC it affected all pairs. This difference at comparatively small distances may be related to the difference of dendrite branching statistics we observed in the topological persistence diagrams of Fig.1E.

These results indicate strong statistical dependence between multiple touches of the same axonal (A) and dendritic (D) trees. A similar observation has been made by Reimann et al. [58], who described statistical dependencies between the formation of a connection from an axon A onto a dendrite D1 and from the same axon onto a different dendrite D2. They demonstrated that such dependencies explain the non-random structure of mouse circuitry and found their source to be the physicality and variability of neurites. While for a class of neurons their axons (or dendrites) reached the entire area all around the soma, each individual one can only cover a small part of the space. That part is different for each neuron, but in a way that is not completely random, as it must be spatially continuous due to the physicality of neurites. This leads to statistical dependencies between connections where the presence of a connection onto neuron N makes the presence of additional connections in the neighborhood of N drastically more likely. We generated 2*D* point clouds (see Methods) to evaluate the spatial properties of the neuronal shapes. Interestingly, while the difference between species in the mean of the dendritic density clouds appeared to be subtle, clear qualitative differences were observed for the axons (Fig. S4). For both axons and dendrites, we observed that a larger area was associated with significant variability for humans than for rat in the CV of the point clouds (Fig.4B). In line with the statistical dependence of connections, stronger variability of axons than of dendrites was found for both human and rat models (Fig.4B).

Reimann et al. [58] quantified the strength of this effect by contrasting the connection probability *P* (*A* → *B*) into a given distance bin with the connection probability conditioned on the nearest neighbor of B being innervated, *P* (*A* → *B* | *A* → *NN* (*B*)) (Fig.4C, top left). To rule out long-tailed degree distributions as alternative explanation, this was contrasted with the same analysis on a configuration model (degree-preserving randomization) of the data. When we performed this analysis on the data of Peng et al. [12], we confirmed that this statistical dependence greatly affects human connectivity (Fig.4C, bottom left). Connection probability for connected nearest neighbors (green) was increased over the base connection probability (black) in almost all distance bins, and this was incompletely captured by the control (dashed lines).

Once again, we emulated a patch-sampling experiment (spatial scale: 120*µm*) in the rat and human CPC networks and repeated this analysis (Fig.4C, center).

While for the human network the we found a positive result at comparable strengths as for the data of Peng et al. [12], for the rat network the effect appeared weak to non-existent. We also performed this analysis for the entire ATC networks (around 20, 000^2^ pairs each). The much larger number of neuron pairs considered gave rise to more robust statistics and allowed analysis of distance bins up to 1000*µm*. We found strong increases of connections probabilities for connected nearest neighbors for both human and rat. However, the magnitude of the increase was larger for human specifically at lower distances, in line with our previous results (see also Fig.S3).

Taken together, the following picture emerges: Local circuitry in neuronal networks is complex in part because the process determining potential connectivity (i.e., touches) is complicated and characterized by multitudes of statistical dependencies within and across pairs of neurons. The specific strength and statistics of the dependencies is determined by the shape and branching of the neurites. In general it is the axon driving most of this effect due to its higher variability across individual neurons within a class (Fig.4B; see also Reimann et al. [58]). However, the source of a significant difference between human and rat networks lies in the branching of dendrites, specifically close to the soma.

## 3 Discussion

In this work, we have explored the effect of neuronal morphologies on the network architecture. We did this by studying the differences between computational models of two species: human and rat, and when possibly verifying the predictions on experimental data. We focused on the network formed by pyramidal neurons in cortical layers 2 and 3, for which more comprehensive human data is available. We have confirmed the results of Kanari et al. [30] that networks predicted from human neurons have higher complexity than networks predicted from rodent neurons. Specifically, we found: Longer-tailed degree distributions, indicating more heterogeneous network connectivity, typically a measurement of network complexity [22, 59, 60]; a stronger effect of spatial scale on connectivity; more specific locations of reciprocal connections; higher motif counts. By also considering the type of algorithm used to predict connections from axo-dendritic touches, and of the spatial scale at which connections are measured, we gained additional insights into why higher network complexity emerges for human.

All together, the following picture emerged. The connectivity backbone was given by the process by which axons meet dendrites, leading to potential connections which are structurally different between human and rat. This difference was reinforced as synapses form for only a small subset of the potential connections in this backbone. Biologically, potential connectivity involves the growth of neurites according to complex developmental signals. Our work did not capture this, but merely approximated the end point of the process by placing reconstructed morphologies into an in silico volume and detecting axo-dendritic touches. This potential connectome was previously described as simply being all-to-all and hence a “tabula rasa” [61]. In contrast, we found it to be very complex with significant statistical dependencies between myriads of neuron pairs and important differences between human and rodent. Naturally, the potential connections (or touches) must be on neurites, and therefore they cover only a small, but spatially continuous part of the volume around the soma. This spatial continuity led to statistical dependencies in two forms: First, each axo-dendritic touch increased the probability that further touches at different locations of the same dendritic tree existed. This led to a distribution of touch count per neuron pair that was more heavy-tailed than expected. Second, an axo-dendritic touch increased the probability that the same axon also touched the dendrites of a different, but nearby neuron. This led to statistical dependencies between connections that explain motif over-expression.

Crucially, the strength and details of the process depend on size, shape and branching of the neurites. In this regard, human dendrites had additional branching near the soma leading to stronger statistical dependencies for neuron pairs within around 100 to 200*µm*. We speculated that this additional branching is possible because of additional space between neurons being available due to the lower cell density in human. This idea had already been formulated by Cragg [27] (see also DeFelipe [1], Cano-Astorga et al. [62]). We must note that for the data of Peng et al. [12] (Fig.4C) we found the strongest statistical dependencies for pairs beyond 100*µm*, arguably contradicting the specific role of pairs at low distances. We suggest that additional data similar to Peng et al. [12] is needed to further resolve this.

The difference between human and rat circuitry was reinforced by the process by which synapses were formed for a small subset of the potential connections. Once again, this is a complex process in biology that our work did not attempt to fully capture. Instead, we formulated statistical constraints on the end point of that process that we derived from experimental data on axon lengths, bouton densities and the number of synapses per connection. In order to fulfill these constraints simultaneously, a connection survival function is needed that depends on the number of axo-dendritic touches for a pair of neurons, in line with the principle of cooperative synapse formation [54]. Here, we found for human a survival function that increased more steeply with touch count, which was supported by the longer neurites simply being able to sustain a more stable number of synapses. This led to a higher likelihood of connections formed by multiple synapses. This would make transmission from the pre-to the post-synaptic neuron more reliable, which is in line with physiological results [5, 9, 10]. Moreover, it implies that for human the presence of a connection is more strongly determined by the touch count of a pair and thus less random.

The structural consequences of all the above are as follows. First, despite longer-reaching neurites, human local connectivity decreased more sharply with distance and was overall more strongly characterized by distance-dependence. This was a consequence of the connection survival function and multiple touches being more likely the closer a pair was. This also led to a larger number of synapses per cell, but lower synaptic density (per unit volume) compared to rodent, which is supported already by the studies of Franz Nissl [28] and Constantin Von Economo [29], as discussed in Cragg [27]. More recently, electron-microscopy images have confirmed larger number of synapses per neuron in human compared to rodent [1], but lower number of synapses per volume unit [1, 4]. Second, overall higher complexity of human circuitry, as measured for example by simplex counts. Third, while the locations of reciprocal connections were non-random in both rat and human, this trend was much stronger in human. Specifically, reciprocity of an edge was structured such that the count of simplex motifs was increased over a random control. We speculate that the last two points can be explained as follows: Because in human the presence of a connection is more strongly determined by touch count, and less random, the complex structure formed by the potential (i.e., touch-based) connectivity is more likely to survive the pruning process and is even enhanced by it.

Our results show that more relevant factors than randomly removing edges or touches from the network could be similar between species [63, 64]. For example, while the overall connection probability of the human and rat ATC networks was drastically different, they reach comparable levels on their corresponding CPC pruned networks (Fig.2E). Thus, our work acknowledges and reinforces the idea that pruning is a crucial process of selecting actual from potential connections, such that the network becomes more robust, efficient and adaptable during development [65, 66]. This provides insight into how synaptic pruning happens in biological systems.

Some of the above has implications that may be unexpected or even controversial. We predict that human circuitry is in some ways more constrained than rat. Lower connection density in ATC, and the stronger decrease of connectivity with distance in CPC, combined with lower neuron densities means that each human neuron selects its synaptic partners from a smaller pool of neurons. Traditionally, we would think of a human as rather the opposite: A being with few innate or pre-determined traits, but greater potentiality, i.e., capability for learning. Although apparently counterintuitive, we note that the greater heterogeneity, longer-tailed degree distributions, and more higher order motifs, only arise due to such constraints. Indeed, they are largely absent in weakly constrained random networks like Erdős–Rényi [40]. Moreover, these features have been found to have positive functional impact: they have been predicted to enhance circuit function [22, 49, 67], and they also ensure greater network functional complexity and heterogeneity that are associated with increased robustness and stability [68, 69]. Additionally, while our results predict stronger constraints on the space of network topologies that can be explored in human circuitry, the space is still vast. At this point it is important to mention that our results are only about local circuitry and for comparing circuits of the same number of neurons. Overall, the much greater number of neurons in the human brain and the strong emphasis on long-range connectivity will more than compensate for this.

Finally, our results also suggest that the amount of overexpression of reciprocal connectivity depends on the spatial scale of sampling, when in vitro-like sampling and analysis methods are employed. As increasingly higher distances are sampled, the distance-dependent connectivity begins to become more important in comparison to Erdős–Rényi models. Experimentalists cannot account for this confounding factor by only having access to low distances, where distance-dependence plays a less significant role. However, such results are not necessarily representative of the entire network’s structure. In the human CPC network, the mean distance of connected pairs was 247*µm*, a scale for which distance-dependence clearly mattered and generated an unexpectedly high number of reciprocally connected pairs.

### 3.1 Caveats and limitations

While we confirmed some of our results in experimentally measured connectivity datasets and our CPC models were validated against experimental data, many of our results still stem from stochastically pruned, predicted connectomes. It has been long discussed to what degree algorithmically such predicted connectivity can make relevant predictions about biological connectivity [36]. In that regard, we suggest that our predictions be further tested in electron-microscopic reconstructions of rodent (mouse, rat) and human connectivity in the future. Specifically, it would be possible to contrast potential (i.e., touch-based) connectivity with actual connectivity in such datasets. In this work, we used electron-microscopy data only for a single supporting analysis for the following reason: Electron-microscopic connectomes require extensive proof reading to provide accurate data. While proof reading has progressed a lot for the MICrONS data, it is not yet at a level sufficient for the analyses in this work for the H01 data.

Furthermore, we want to stress once more that we do not claim that placement of reconstructed morphologies in an *in silico* volume fully captures the developmental processes that give rise to potential connectivity. Nor that the algorithm of Reimann et al. [32] underlying the CPC networks captures all aspects of synapse pruning. We only state that they capture the statistics of the outcomes of those processes. This is indicated by them being validated against experimental data in the literature [34] and in this work. Hence, we can use them to make the statistical predictions listed above.

Importantly, the literature provides a number of examples where such statistical predictions are not valid. Reimann et al. [34] found that targeting of inhibitory classes described in Schneider-Mizell et al. [70] is reproduced only if specific biases are applied during pruning. The result does not affect this work, as we focus on excitatory connectivity only, where targeting appears less specific [71]. Moreover, Motta et al. [33] showed that synaptic targeting of subcellular domains is not reproduced when pruning is statistically independent with uniform probabilities. This result does not affect this work either, as it only investigates pruning akin to our T_to_T algorithm; whereas our cellular predictions result from CPC networks, which Reimann et al. [34] have shown to recreate the apparent targeting described in [33]. We further expand the negative results for T_to_T pruning, a common interpretation of Peters’ rule, as the results of this method were furthest from the experimental results of [12]. Furthermore, connectivity predictions at cellular resolution from potential connectivity have also been successful in the past. Cook et al., [72] have shown that such an approach can predict the C. elegans connectome.

Additionally, all the differences between human and rat we described might instead be attributed to other factors such as brain region, subject age, or data sparsity rather than inherent differences between species. In particular, the data used to build the models belonged to different cortical areas that process different types of information: temporal cortex (Ta21 or Ta22) for human and primary somatosensory cortex for rat. However, the anatomical differences we considered such as: cell density, ratios of excitatory versus inhibitory cells and neuronal morphological characteristics, have also been observed in clearly comparable brain areas, such as the hippocampus, between human and rodents. e.i., Human hippocampal CA1 pyramidal neurons have more complex branching around the soma compared to the corresponding mouse hippocampal pyramidal neurons [30, 73, 74]. Moreover, within the same species, it is known that the complexity of the neuronal apical branch is correlated to the layer thickness in the cortex [75]. Consequently, neuronal morphologies belonging to cortical areas with similar layer thicknesses should be comparable within the same species. This is the case for the rodent somatosensory and temporal areas and we extended this assumption to human.

With respect to age ranges, for rat we used morphologies from juvenile subjects (p14 - p16), while for human we gathered pyramidal cell morphologies from subjects of very different ages (19 - 85 years old). Although the brain suffers drastic changes during development [76], a recent study comparing excitatory neurons in human layers 2 and 3, from subjects with ages between 0 to 73 years old, found no significant differences regarding some morphological features such as: dendritic length and total number of dendritic branching. The biggest morphological and physiological differences were observed between children (0 - 12 years old) and adults (20 - 85 years old) [77], which supports the morphological and anatomical comparisons made in this study.

## 4 Methods

### 4.1 Biologically detailed cortical models

The models used for this study are biologically detail models of the rat somatosensory cortex and the human temporal cortex. The rat model was published in 2015 by Henry Markram’s group [17] and it has been recently updated [34]. In that publication they presented the development of a circuit building workflow and tools for each of the pipeline steps. The same approach was taken to produce the human cortical model [11], although some adaptations had to be applied.

In brief, the human cortical model has been originated by collecting data on dendro-axonic morphological reconstructions, anatomy, cell density and composition per cortical layer, as well as data on bouton density per cell type and the number of synapses per connection between cell pairs. Some of the data were collected from the literature. All model details have been published in Barros et al. [11].

### 4.2 Connectomes and network types

#### 4.2.1 The “All touches connectome” (ATC)

To construct the circuits for human and rat, biological data on cell placement, cell density and cell composition per layer were used as described in Markram et al. [17], Reimann et al. [34]. Then, 656 for human and 1191 for rat unique reconstructed morphologies were placed in the volume following the workflow of [34] for all layers. The *All touches connectomes* were built from this data by considering all the potential connections as the axo-dendritic and dendro-dendritic appositions between pairs of neurons within a distance of 2 *µ*m as actual “contact” points (see results section of [11] (human) and [17] (rat) for more details).

In order to reduce the computational costs, we extracted for each of the networks a central subvolume of approximately 20000 neurons (20072 for human, 20064 for rat). We represent any network, with nodes (neurons) and an edge from one neuron to another if there is at least one touch or contact point in that direction (Fig.2A, top left). Consequently, it can be represented by a non-symetric matrix *M* where *M*_*ij*_ is the number of touches from neuron *i* to neuron *j*. In particular, the total number of connections is the number of non-zero entries of *M* and the total number of touches is the sum of all the entries of *M*.

#### 4.2.2 Human control using rat data (HC_RD)

This model aims to answer what connectivity would be expected if rat connectivity was adjusted to the overall dimensions found in human. In particular, it considers the following high-level differences, observed for excitatory neurons in cortical layers 2/3, between the two organisms [11]:

- The volumetric density of neurons is higher in rat than in human. For this, we use factor *h* = 2.8
- As a consequence, human neurons are further apart from each other. For this, we use factor 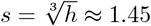
- The mean total dendritic length is longer in human than in rat by a factor of *r*_*dend*_ = 2.75
- The mean total axon length is shorter in human than in rat by a factor of *r*_*axon*_ = 0.7
- The bouton density, and hence the density of efferent synapses per unit of axon length is unchanged between the organisms: *b* = 1.

We note that the combination of values for *h* and *r*_*dend*_ results in almost the same *volumetric* length density of dendrites between rat and human: There are fewer neurons in a given volume for human, but each individual one compensates by being larger. This is in line with a similar observation for mouse and human in Kanari et al. [30].

We begin with a thought experiment: What if a rat microcircuit was merely “stretched out” to human dimensions? This could be conceptually performed by simply compressing the spatial axes, such that, e.g., 1*µm* only represents 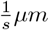. As this is merely a re-labelling of axes, the potential synapses between neurons would remain identical. However, expressed as a distance-dependent connection probability the distance scaling would have to be applied. For example, in the case of exponentially-decreasing connectivity:

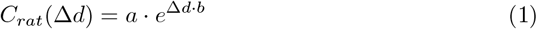

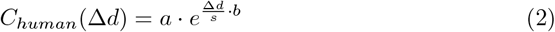

While this takes into account the thinning of neuron densities, this would result in the total axon and dendrite lengths being scaled by the same factor *s* instead of the actual values. To take the actual values into account, we assume that the extra dendrite (and loss of axon) is distributed proportionally over all distances. Specifically, we assume that the maximum spatial reach of dendrites/axon, and hence the distance-dependence of connectivity remain unchanged. In that case, connectivity would simply scale with the extra/loss of neurites:

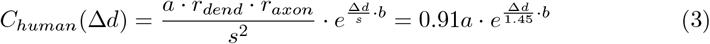

The instances of this model are then constructed as follows. First, for the CPC network of rat, we fit exponentially decreasing probability profiles (as in Equation 1) for the existence of a connections with at least one, two, three, etc. potential synapses. Explicitly building models for successively higher number of synapses takes into account the effect outlined in Fig.4A, right. These profiles are then “stretched” and scaled as in Equation 3. The result is a stochastic control for the number of potential synapses between pairs of neurons expected for human data based on high-level anatomical differences. We evaluate it to generate any number of stochastic instances of it. As we constructed distributions one, two, three, etc. potential synapses, the instances not only prescribe the locations of connections, but also the number of potential synapses forming them. As such, the instances can be subjected to the pruning methods described in the previous sections.

#### 4.2.3 Pruning methods

The ATC and HC_RD networks were *pruned* i.e, touches or edges were removed, to reach biologically realistic connectivity. Several pruning methods were used; they are summarized in Table S1 and visually represented in Fig.2A.

First, the pruning algorithm of Reimann et al., generating what we called the *Cooperatively Pruned Connectome* (CPC), as it employs the principle of cooperative synapse formation [32]. Second, four pruning methods using CPC as a reference which fall in two broad categories, which we describe below.

##### Pruning to match touches

Given an ATC network represented by a matrix *M* we construct a pruned network that has the same number of touches than its corresponding CPC network *M*_*CP C*_. We do this in two ways:

1. Pruning touches at random, which we denote by *T_to_T*, to represent pruning touches to match touches.
2. Pruning edges at random, which we denote by *E_to_T*, to represent pruning edges to match touches. We do this heuristically via an iteratively procedure until the difference between the pruned matrix and *M*_*CPC*_ is at most 1% of the touches.

##### Pruning to match edges

Given an ATC network represented by a matrix *M* we construct a pruned network that has the same number of edges than its corresponding CPC network *M*_*CPC*_. We do this in two ways:

1. Pruning edges at random, which we denote by *E_to_E*, to represent pruning edges to match edges.
2. Pruning touches at random, which we denote by *T_to_E*, to represent pruning touches to match edges. We do this iteratively until the difference between the pruned matrix and *M*_*CP C*_ is at most 1% of the edges. Note that this is equivalent to pruning edges, where the probability to keep an edge is proportional to the number of touches it has.

#### 4.2.4 Other control networks

In addition to networks obtained from pruning the ATC or HC_RD networks using the CPC as reference, we also generated stochastic networks that preserve specific structural properties of the CPC networks. All these were generated using the using the `Connectome-Analysis` package [22].

- The *Erdős–Rényi* controls (ER) are random directed networks, where each edge is added independently at random with a fixed probability *p* given by 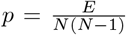, where *E* and *N* are the number of edges and nodes in the original network, thus keeping the global connection probability fixed.
- The *directed configuration model* controls (CM) are random networks with (approximately) fixed in- and out-degree sequences, i.e., the vectors whose entries are the in- and out-degrees of all the nodes in the network.
- The *distance dependent model* controls (DD) are random networks where the edges are added independently at random with a probability that is exponentially decreasing with distance. That is, for any pair of neurons at Euclidean distance *d*, their probability of connection is given by

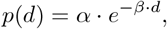

where *α* (the probability at distance zero) and *β* (the decay constant) are determined from the connectivity of the original graph.
- Finally, the *bishuffled* control. To build such a control, for every reciprocal edge in the original network we chose one direction to delete at random. Then we introduced back at random the same number of reciprocal connections along the existing edges.

### 4.3 Network metrics

#### 4.3.1 Connection probability

We consider different types of connection probabilities a global network metrics and we describe them now in order of complexity.

The *global connection probability* of a network, is the overall probability of two neurons to share a connection in either direction. Given an *N* × *N* connectivity matrix *M* it is given by 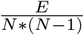 where *E* is the number of edges of *M* i.e., the number of non-zero entries.

The *connection probability per mtype-pathway*, is the overall probability of a neuron in mtype *A* to have a connection to a neuron of mtype *B*. If *A* and *B* are the same mtype, then the connection probability is compute as above but restricted to the submatrix of the given pathway. If *A* ≠ *B*, then this connection probability is given by 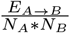, where *E*_*A*→*B*_ is the number of connections from mtype *A* to mtype *B* and *N*_*A*_, *N*_*B*_ are the number of neurons of mtype *A* and *B* respectively.

Finally, we consider as well the *connection probability at a given distance*. For this, we bin all possible distances between all pairs of neurons in *M* (bin sizes: 50 *mu*m for rat, 50*1.76 *mu*m for human). For each distance bin *D*, there is a subset of all pairs of neurons at that given distance. Thus the connection probability at distance *D*, is given by the number of connections between such pairs divided by the number of total pairs. We model the decay of the probability of connectivity with an exponential decay function *P* (*d*) = *ae*^−*bd*^, where *P* (*d*) is the probability of connection at distance *d* and *a, b* are real numbers. We implement this using the `Connectome-Analysis` python package [22].

#### 4.3.2 Survival probability of an edge

Beyond comparing the connection probabilities of different pruned networks, we also examined the identities of the edges that were removed. To this end, we generated 30 independent instantiations for each pruning type. We evaluated the probability of *survival of an edge* i.e., the likelihood that a given edge is retained after pruning, as a function of its number of touches. To compute this, we grouped edges in the CPC network by their number of touches and, for each instantiation, calculated the fraction of edges in each group that “survive” the pruning process. This procedure was applied across all instantiations and pruning methods.

#### 4.3.3 Formation of subsequent touches

While the connection probability described in the previous section considers only the existence of at least touch (or synapse) between pairs of neurons, we also analyzed the formation of additional ones beyond the first. We study the number of touches of connected pairs at a given distance, using a model that is consistent with exponential decay connection probability *P* (*d*) defined in the previous section.

For this, we bin all possible distances between all connected pairs of neurons in *M* (bin size: 50*µm*). In each distance bin *D*, and for each number of touches *t >*= 1 we count the number of connected pairs in that bin that have at least *t* touches, which we denote by *C*_≥*t*_ (*D*). Additionally, we define *C*_≥0_(*D*) as the total number of pairs in that bin.

We then compute the function

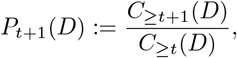

which computes the probability that a pair at distance *D* has more than *t* touches given that it was already *t* touches. To ensure robustness, we exclude distance bins where *C*_≥*t*_ (*D*) *<* 500. We study this function for 1 ≤ *t* ≤ 10. The function *P*_*t*_(*D*) can be thought of as a refinement of the connection probability function *P* (*D*). Note in particular that *P*_1_(*D*) is equivalent to the connection probability.

#### 4.3.4 Degree

We study the indegree, number of neurons in the pre-synapatic population, and outdegree, number of neurons in the post-synaptic population for the E_to_E network. To evaluate its heterogeneity, we measure the *skewness* of the distributions using `SciPy` [78]. The skewness of a unimodal distribution measures the direction and relative magnitude it deviates from a normal distribution. The skewness of a normal distribution is 0, a skewness value greater smaller than zero means a right respectively left tailed distribution.

#### 4.3.5 Motif counts

To bridge local and global network structures we count several type of motifs. *Reciprocal connections* or bidirectional edges, are pairs of neurons synaptically connected in both directions. We count this for the full circuits as well as for simulations of patch-clamp experiments at different scales. To determine over and under expression of these motifs, we normalized them with respect to random controls of the same size: ER controls for the patch clamps and ER and DD controls for the full circuits.

*Triad motifs* are subgraphs on three nodes. There are 16 types of directed graphs on three nodes up to isomorphism (i.e., up to a relabeling of the nodes) see (Fig.S2E). We count only connected triad motifs, of which there are 13 using the `Connectome-Analysis` package [22]. For the in-silico patch clamp experiments, we count triads exactly. For the full ATC networks, we approximate the total counts by classifying at most 5000000 triads sampled at random and linearly extrapolating the total counts. To determine over and under expression of these motifs, we normalized them with respect to random controls of the same size: ER controls for the patch clamps and ER and DD controls for the full circuits.

A *directed k-simplex* is a motif on *k* + 1 neurons all to all connected in feedforward fashion. That is, there is a numbering of the nodes 0, 1, … *k*, such that there is an edge from *i* to *j* whenever *i < j*. Edges in the other direction are allowed to exist. In particular, 0-simplices are the nodes of the network, 1-simplices are the edges of the network and 2-simplices are transitive triads (see Fig.S2F for examples). We counted directed simplices using the `Connectome-Analysis` python package [22].

### 4.4 Analysis of statistical dependencies of connectivity

We extend the concepts of connection probability and non-independence of connectivity at a given scalar distance to include offsets in two dimensions, accounting for variations in both the vertical direction and the horizontal plane. We studied the rat and human in-silico models as well as the mouse connectome provided by MICrONS [57]. Given that we will just consider the presence or absence of a connection and not the number of touches, we represented the connectomes as binary adjacency matrices.

For each connectome, we calculated for each pair of neurons their offsets in the horizontal plane and along the vertical axis. Offsets were then binned in both directions with a bin size of: 50*µm* for rat or mouse connectomes and 1.76 · 50*µm* for human connectomes. The 1.76 scalar was chosen as the ratio between the nearest neighbor distance of neurons in human data divided by the same in rat data. Let *B*(*a, b*) be the binning function that associates a pair of nodes with a bin. One can think of a bin as a discretized 2D-vector that describes the offset from the position of *a* to that of *b*. Thus, we refer to a generic bin as 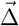.

We computed the outgoing and incoming connection probabilities at a given offset bin over the entire connectome. That is,

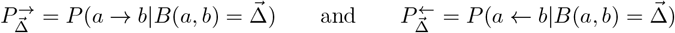

respectively, where the arrow represents the presence of an edge in the indicated direction. This is a natural extension of the connection probability at a distance to two dimensions and to include vertical direction. The first (outgoing) measures the probability of connection in the same direction as 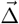, while the second (incoming) measures it in the opposing direction.

Let *NN* (*a*) indicate function that assigns to a node *a* its *nearest neighbor* according to its spatial location. We calculated the connection probabilities of each bin, conditioned on the nearest neighbor of a neuron being connected in the given offset. That is,

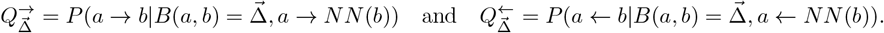

Note in particular, that given our bin size choice, most of the time *a* and *NN* (*a*) belong to the same spatial bin. In this case, 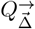 is the probability of more than one connection existing in the offset bin 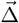, given that one already exists. Therefore, the difference between 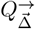 and 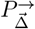 serves as a measure of the degree to which the connectivity in a given bin is statistically non-independent.

### 4.5 Morphological Analysis

#### 4.5.1 Density Clouds

To characterize the spatial organization of neurites, we generated density cloud–based representations inspired by prior approaches for spatial point pattern analysis and neuronal morphology quantification [79, 80]. Traced neurite skeletons were used as inputs to obtain the original point clouds, which were subsequently normalized within each dataset and imaging modality to a common coordinate range (neurons starting at (0, 0)). From these normalized point clouds, we constructed two-dimensional density maps, which are termed density clouds. For 2*D* density maps, points of each neuron were randomly rotated around the Y-axis (which is aligned with the apical dendrites), projected onto the X-Y plane and discretized into bins spanning the interval X: [-1000 × 1000], Y: [-1500, 1500]. Axon and dendrite masses were projected separately onto the X–Y plane.

For the 2*D* density clouds, we computed the mean and coefficient of variation (i.e., standard deviation over mean, denoted CV) of the density clouds across neuron morphologies (Fig 4, Fig S4). The CV of the density clouds across neuron morphologies measures this variability in a normalized way. Sample sizes were: 366 and 103 dendritic density clouds and 78 and 68 axonal density clouds for human and rat respectively. To reduce sampling noise and emphasize mesoscale structure, all histograms were smoothed by convolution with a Gaussian kernel. Bin sizes used matched those of Figure S3 (50*µm* for rat and 50 * *s* = 88*µm* for human).

**Figure 4.**
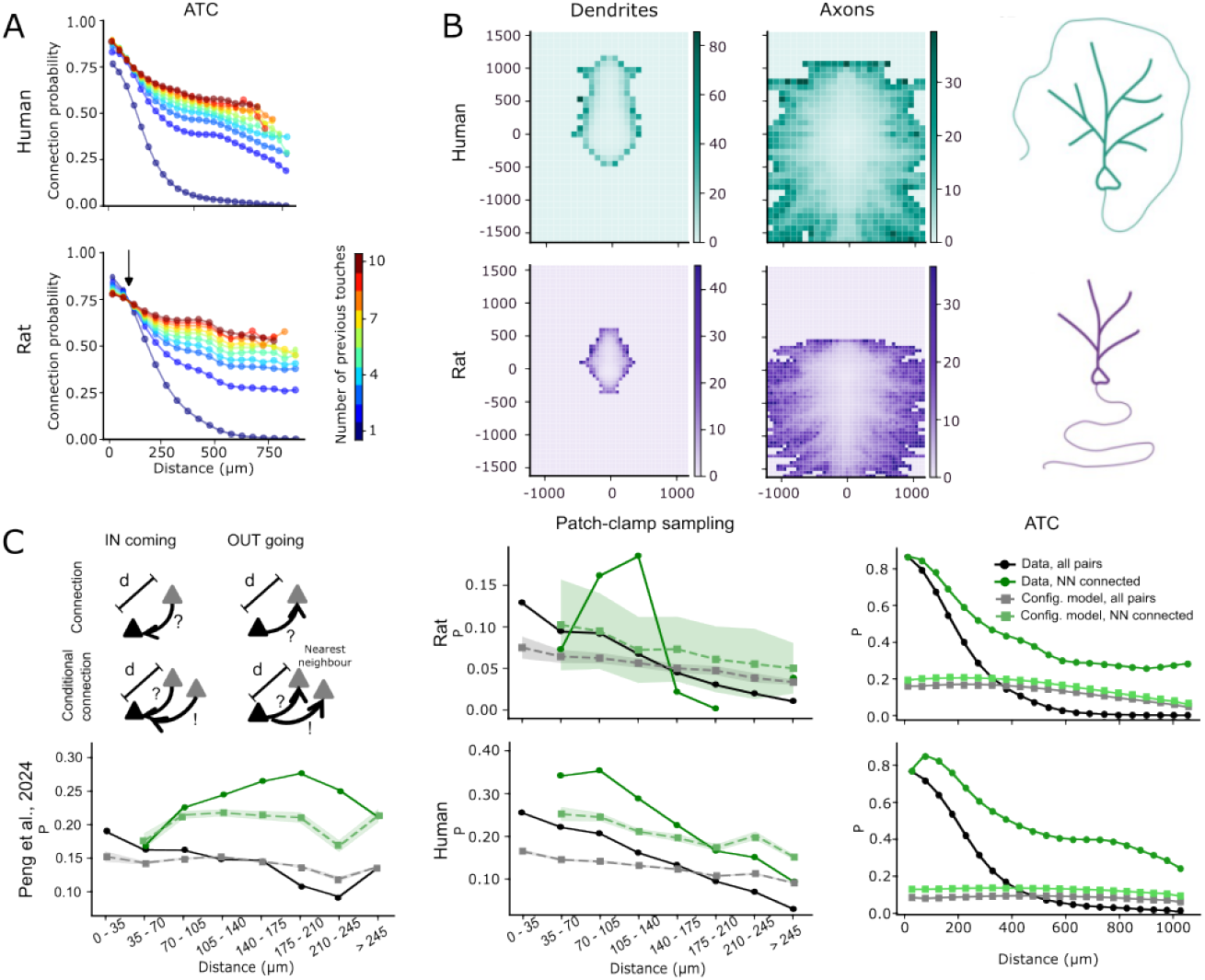
Nearest neighbor connectivity and the spatial orientation of morphologies explain network complexity. (A) Connection probabilities along distance considering the existence of previous touches between pairs, computed for the human (top) and the rat (bottom) ATCs. The color gradient represents the number of previous touches from 0 (dark blue) to 10 (dark red). The arrow points to the shift in connection probability along distance at 100 µm for the rat ATC. (B) Morphological analysis of human (top, teal) versus rat (bottom, purple) pyramidal cells. The morphologies of dendrites (and axons) were randomly rotated (20 times) and projected to x-y plane to generate a density cloud of each neuron. The standard deviation over the mean density cloud for the population of (366 human and 103 rat) dendrites (left) and (78 human and 68 rat) axons (middle). Schematic representation of pyramidal cells branching for rat (bottom right) and human (top right). Thin branches represent axons, thick branches are dendrites. (C) Schematic showing the measured effect (top, left): We contrast the connection probability into a distance bin (top) with the connection probability conditioned on the nearest neighbor of a neuron being connected (bottom). If it affects connectivity, the conditional probability will be increased. (bottom, left) Results of the test in C-top-left for outgoing connectivity applied to the data of Peng et al. [12] Black: overall connection probability in the data; green: conditioned on the nearest neighbor within a patch-sample being connected. Grey and light green indicate corresponding results for a configuration-model, i.e., a control that captures the effect of long-tailed degree distributions. Dashed lines shows the mean and shaded area the standard error of the mean over 1000 instances. (C, middle) As C-left-bottom, but for the data of a simulated patch-sample experiment using the rat (top) and human (bottom) CPC networks.(C, right) As C-left-bottom, but for the the rat (top) and human (bottom) ATC networks.

#### 4.5.2 Topological Morphology Descriptor

Persistent homology [81] captures the topological properties of shapes and has been used for a variety of tasks, including the reconstruction, recognition, and matching of objects. Persistence homology captures relevant information on the underlying shapes by pairing critical values of a function, represented as points in a 2D diagram, the persistence diagram. Typically, the function used as a filtration in persistence homology is the height function [81–83]. In Kanari et al. [31] we introduced an alternative algorithm to compute persistence homology by generating a topological descriptor of neurons (Topological Morphology Descriptor, TMD), which produces a persistence barcode from any tree-like structure (Fig.S5)

The topological morphology descriptor (TMD) [31, 51] transforms tree-like structures, such as neuronal morphologies, with a real-valued function *f* : *V* → ℝ on the tree nodes, to a persistence barcode through a filtration function. It produces an embedding of the graph in a two-dimensional space that has been shown to capture well the topological properties of the graph and has been used for the classification and clustering of rodent [51] and human [30, 39] pyramidal cells. Each branch within the tree is represented by a line in the barcode, which encodes the first and the last time (in units of the function *f*) that the branch was detected in the tree structure. The persistence barcode is a collection of lines that encode the lifetime of each branch in the underlying tree structure. Alternatively, the start and end times of the branches can be represented as two-dimensional points in a persistence diagram. The two representations are equivalent and will be used interchangeably in the manuscript.

## Supplementary material

**Table S1.**
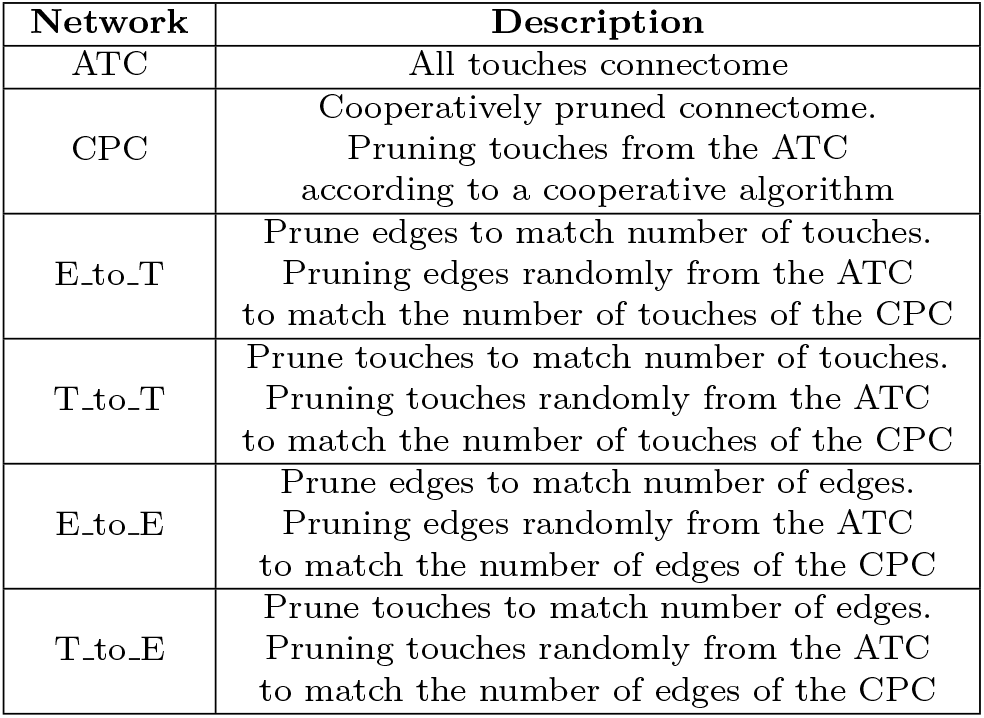
Description of the acronyms used to define the different networks.

**Figure S1.**
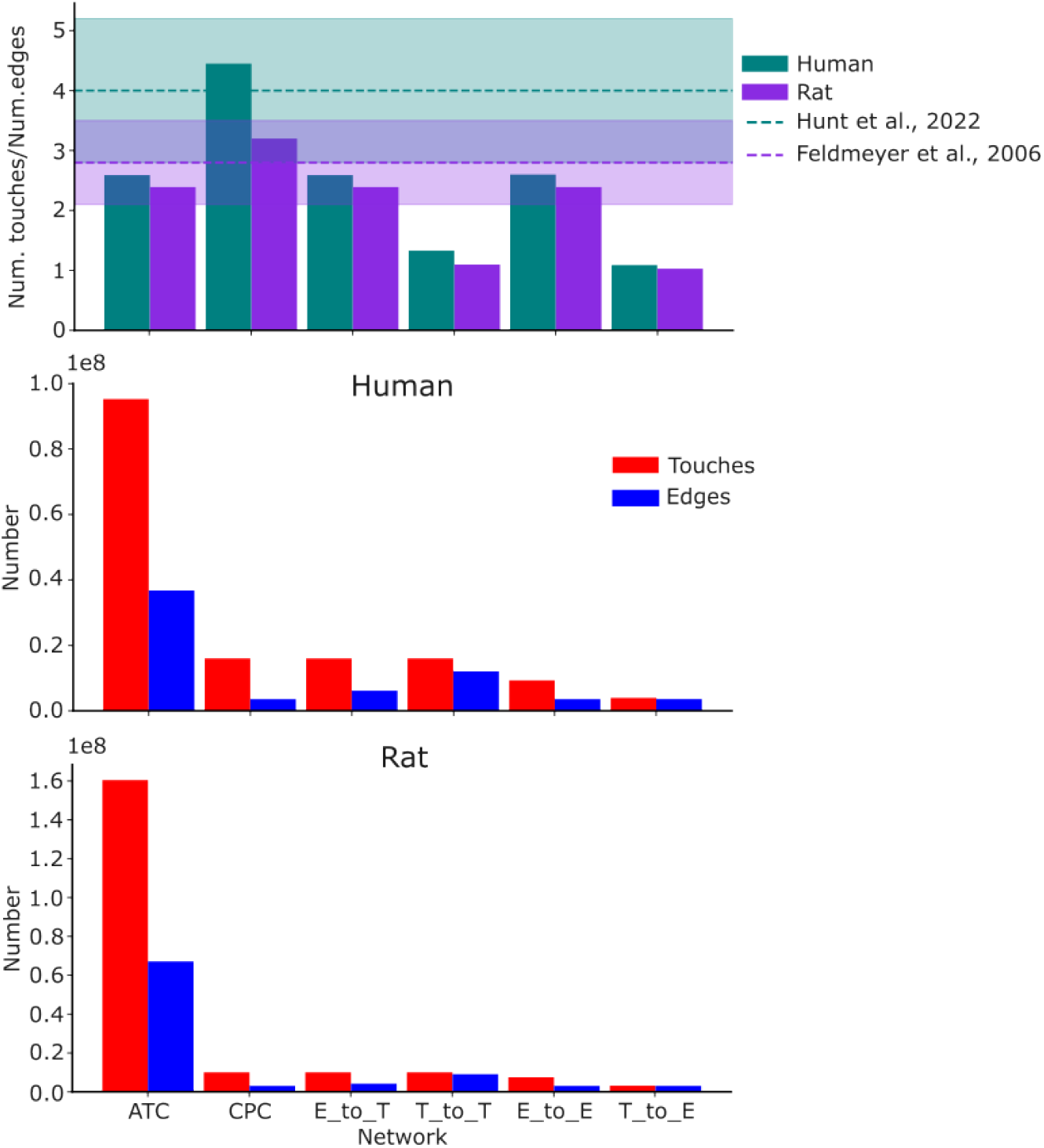
Rates of number of touches over the number of edges for each network for human (teal) and rat (purple). Dashed lines show mean value references from literature and shadows are the standard deviations respectively (top). Total number of touches (red) and edges (blue) in each built network, for the human (middle) and rat (bottom)

**Table S2.**
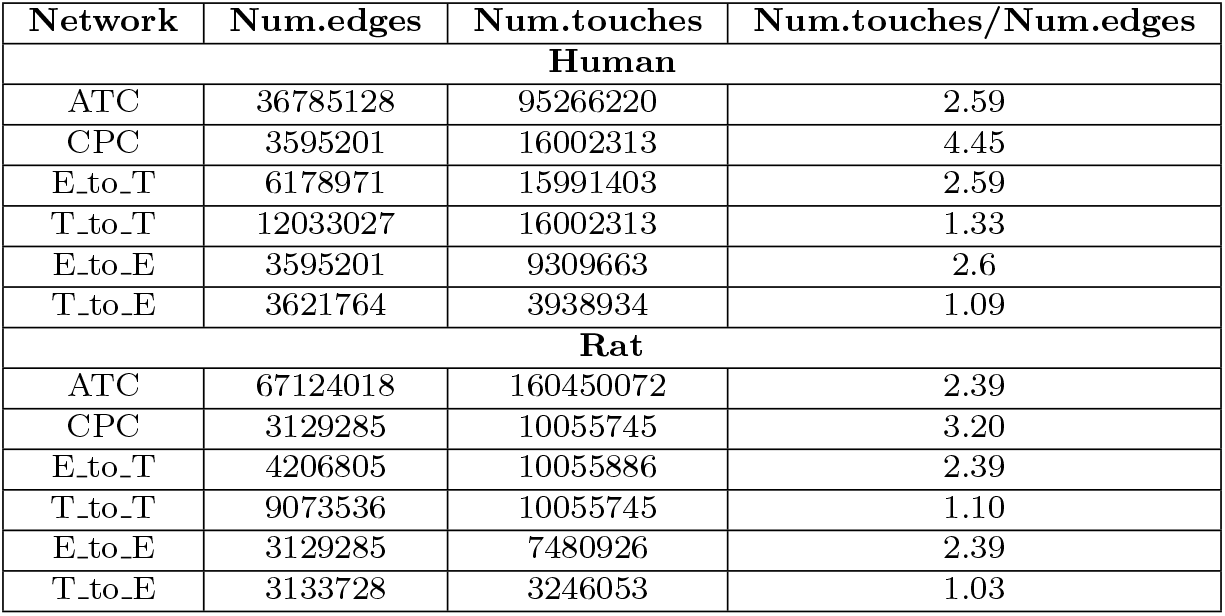
Total numbers of touches and edges and their rates (num.touches over num.edges) for all the built connectomes.

**Figure S2.**
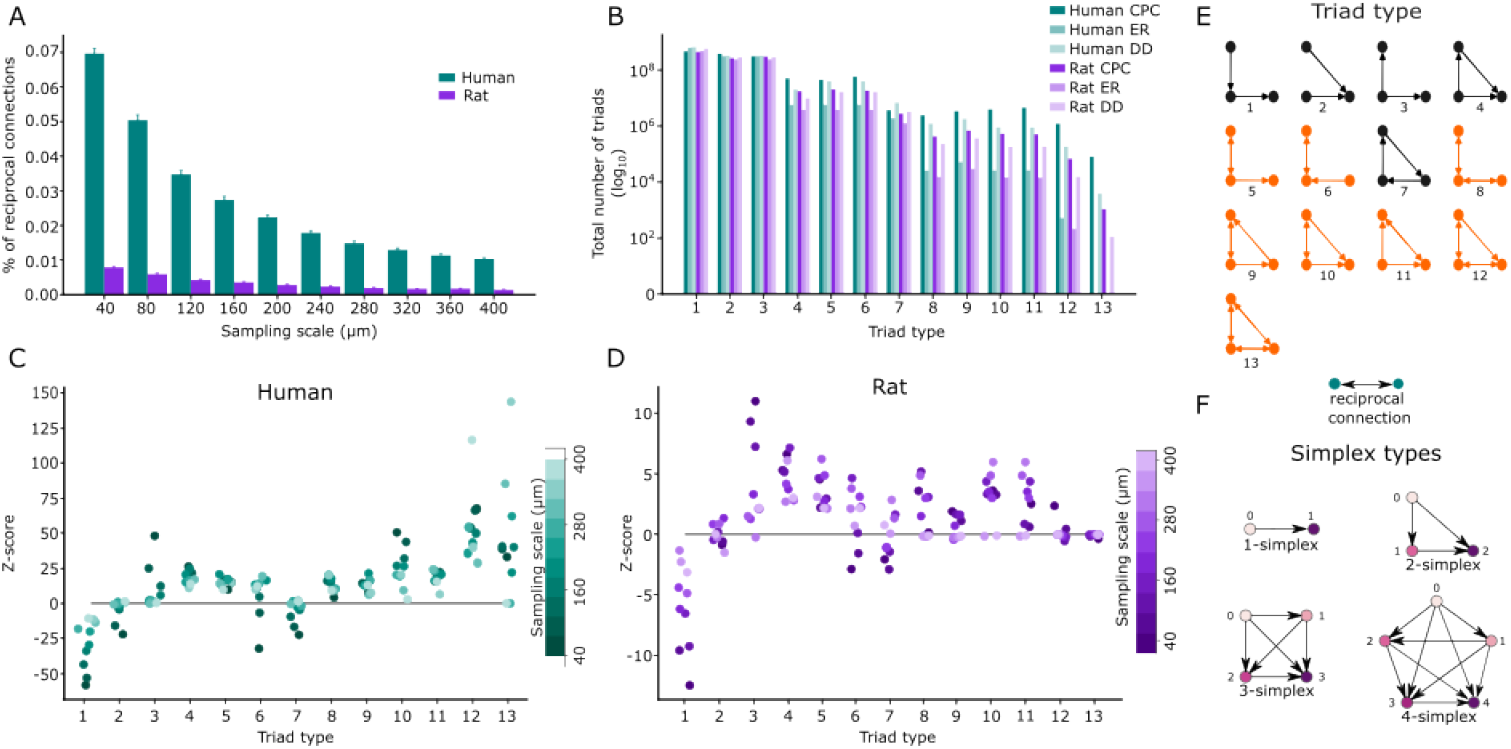
(A) Bar plot with the percentages of reciprocal connections at different sampling scales from (40 - 400) µm for the CPCs of human (teal) and rat (purple). (B) Total number of triads for CPC networks (darker colors) and two random models: Erdos-Reyni (ER) (light colors) and distance-dependent (DD) (lighter colors). (C) Z-score of triad motives expression at different scales (40 - 400 µm for the human CPC network. (D) Same as C for the rat CPC network. (E) Cartoon describing the types of triad motives explored. In orange, triad motives containing reciprocal connections. (F) Cartoon explaining the types of simplices.

**Figure S3.**
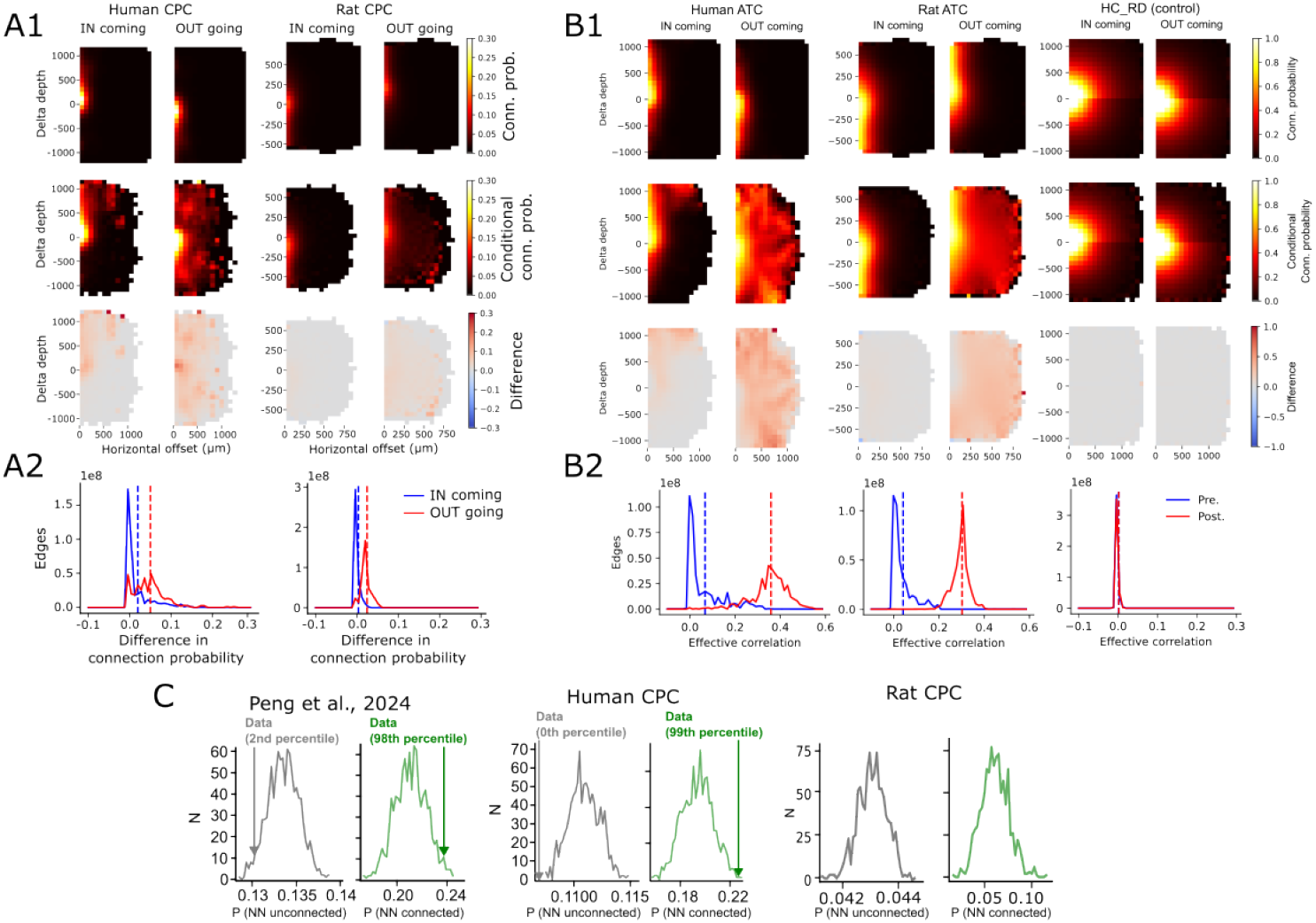
(A1) Contrast between connection probabilities into a distance bin, with the connection probability conditioned on the nearest neighbor of a neuron being connected. All pairs were considered. Additionally, two-dimensional offset bins are considered instead of distances in one dimension. Overall connection probability in the indicated direction (top),conditioned on the nearest neighbor being connected (middle), difference (bottom). Strength of the effect captured in human (left) and rat (right) CPCs. (A2) Distribution of the difference of connection probability across all edges of the corresponding network, incoming (blue) and outgoing (red) connectivity. Dashed lines indicate respective means. (B1 and B2) Same as A1 and A2, respectively, for ATC networks of human (left), rat (middle) and the HC_RD (right). (C) Distribution of overall and conditional connection probabilities at all distances for 1000 configuration model controls of the indicated networks. The arrows indicate the location of the corresponding value in the actual (non-control) networks.

**Figure S4.**
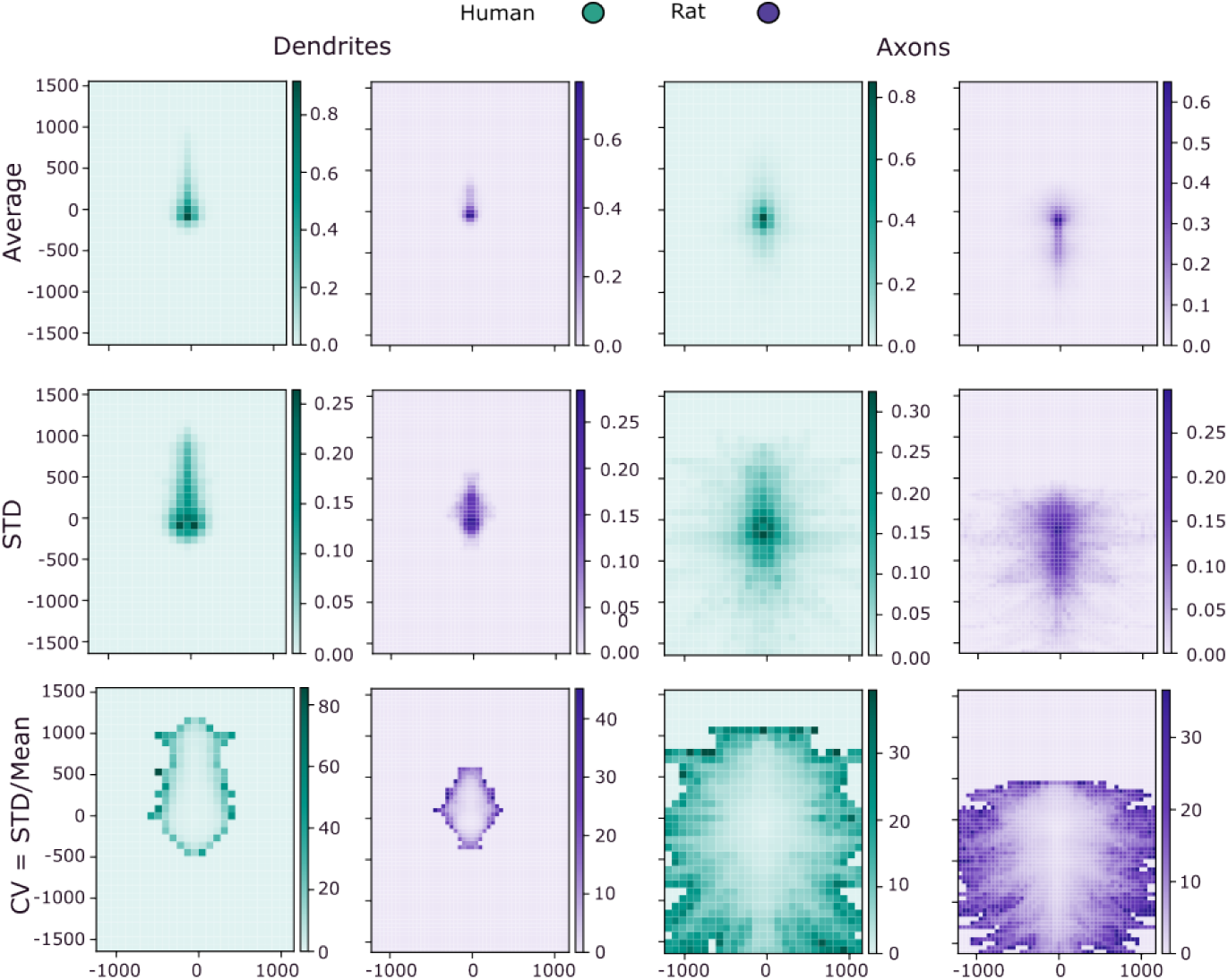
Density clouds. Dendrites and axons were randomly rotated (20 times) and projected to x-y plane to generate a density cloud of each neuron. From this it was computed: the Average (top), standard deviation (middle) and coefficient of variation (bottom) for dendrites (left) and axons (right)

**Figure S5.**
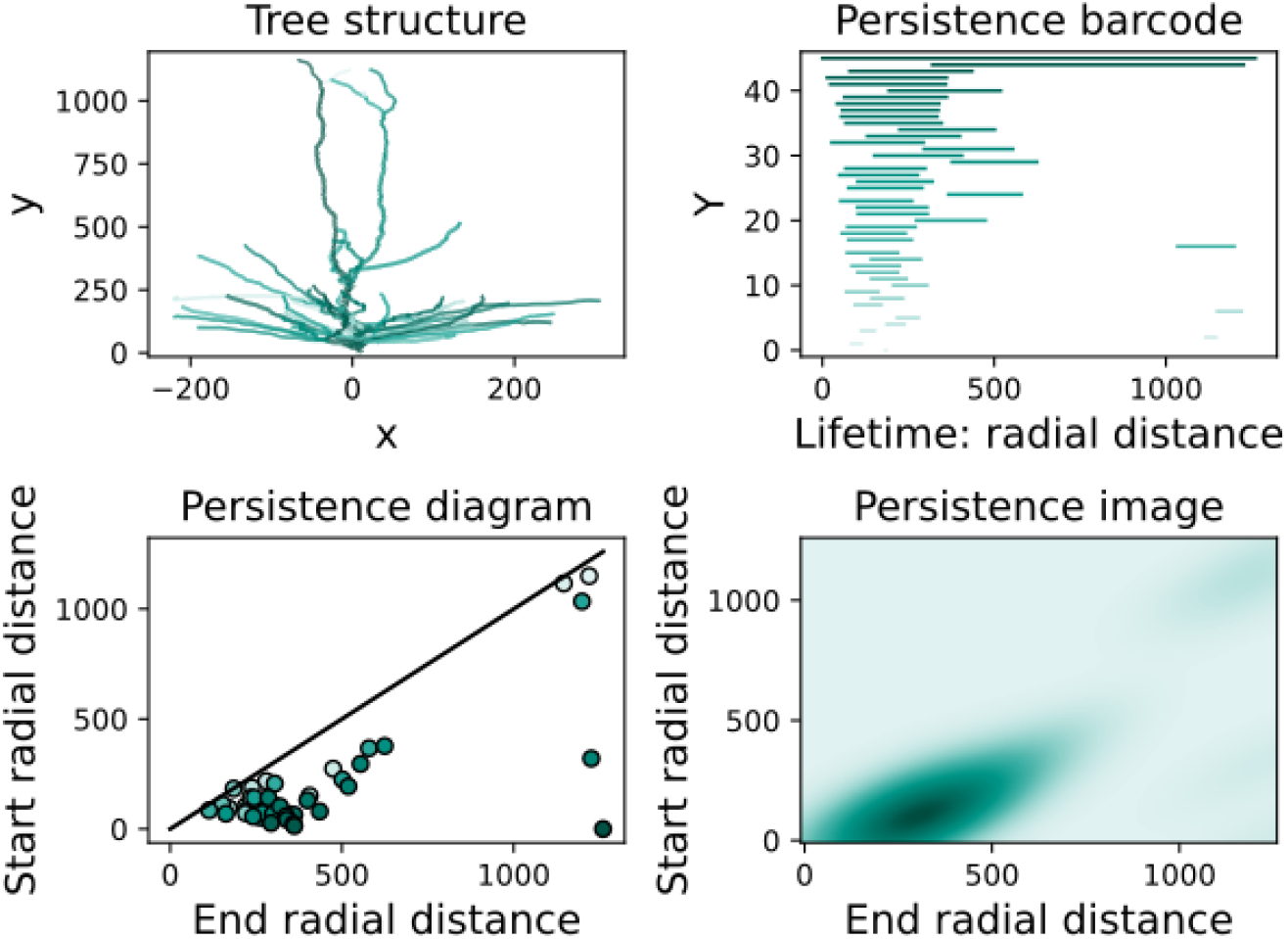
Dendritic tree structure of an exemplar of a pyramidal neuron (top left). Persistent analysis from Kanari et al. [31]: bar-code (top right), diagram (bottom left) and image (bottom right).

