## Supplemental table and figures for "Uncovering the Basis of Human Connectome Complexity: The Role of Neuronal Morphology"

### Supplementary material

**Table S1** Description of the acronyms used to define the different networks

| Network | Description |
|---|---|
| ATC | All touches connectome |
| CPC | Cooperatively pruned connectome. Pruning touches from the ATC according to a cooperative algorithm |
| E_to_T | Prune edges to match number of touches. Pruning edges randomly from the ATC to match the number of touches of the CPC |
| T_to_T | Prune touches to match number of touches. Pruning touches randomly from the ATC to match the number of touches of the CPC |
| E_to_E | Prune edges to match number of edges. Pruning edges randomly from the ATC to match the number of edges of the CPC |
| T_to_E | Prune touches to match number of edges. Pruning touches randomly from the ATC to match the number of edges of the CPC |

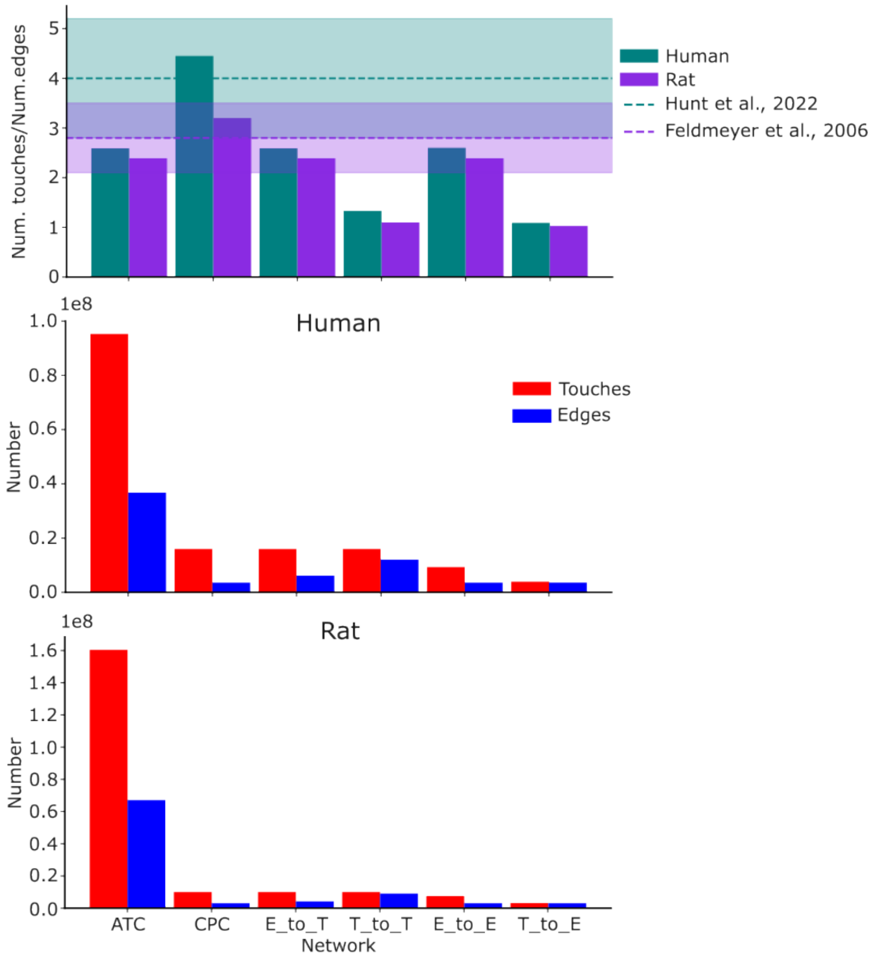


**Figure S1** Rates of number of touches over the number of edges for each network for human (teal) and rat (purple). Dashed lines show mean value references from literature and shadows are the standard deviations respectively (top). Total number of touches (red) and edges (blue) in each built network, for the human (middle) and rat (bottom)

**Table S2** Total numbers of touches and edges and their rates (num.touches over num.edges) for all the built connectomes.

| Network | Num.edges | Num.touches | Num.touches/Num.edges |
|---|---|---|---|
| **Human** | | | |
| ATC | 36785128 | 95266220 | 2.59 |
| CPC | 3595201 | 16002313 | 4.45 |
| E_to_T | 6178971 | 15991403 | 2.59 |
| T_to_T | 12033027 | 16002313 | 1.33 |
| E_to_E | 3595201 | 9309663 | 2.6 |
| T_to_E | 3621764 | 3938934 | 1.09 |
| **Rat** | | | |
| ATC | 67124018 | 160450072 | 2.39 |
| CPC | 3129285 | 10055745 | 3.20 |
| E_to_T | 4206805 | 10055886 | 2.39 |
| T_to_T | 9073536 | 10055745 | 1.10 |
| E_to_E | 3129285 | 7480926 | 2.39 |
| T_to_E | 3133728 | 3246053 | 1.03 |

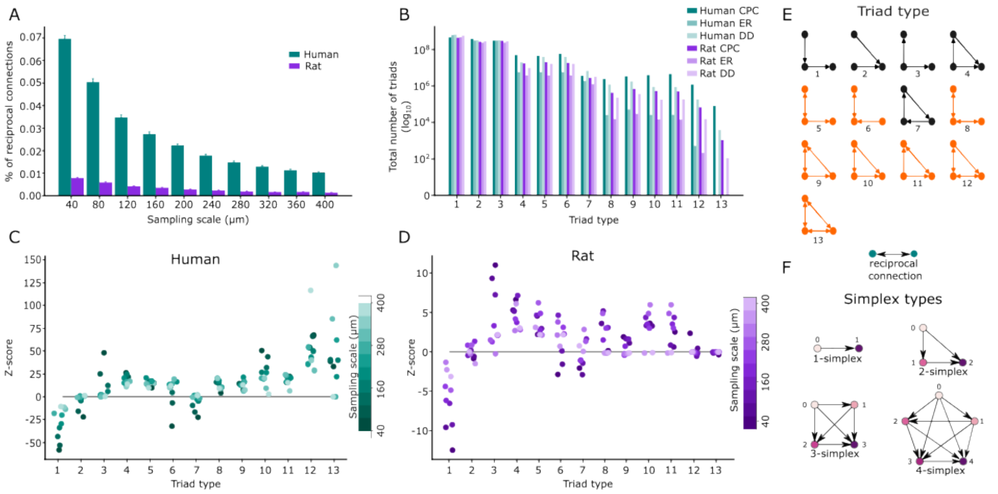


**Figure S2** (A) Bar plot with the percentages of reciprocal connections at different sampling scales from (40 - 400) µm for the CPCs of human (teal) and rat (purple). (B) Total number of triads for CPC networks (darker colors) and two random models: Erdos-Reyni (ER) (light colors) and distance-dependent (DD) (lighter colors). (C) Z-score of triad motives expression at different scales (40 - 400 µm) for the human CPC network. (D) Same as C for the rat CPC network. (E) Cartoon describing the types of triad motives explored. In orange, triad motives containing reciprocal connections. (F) Cartoon explaining the types of simplices.

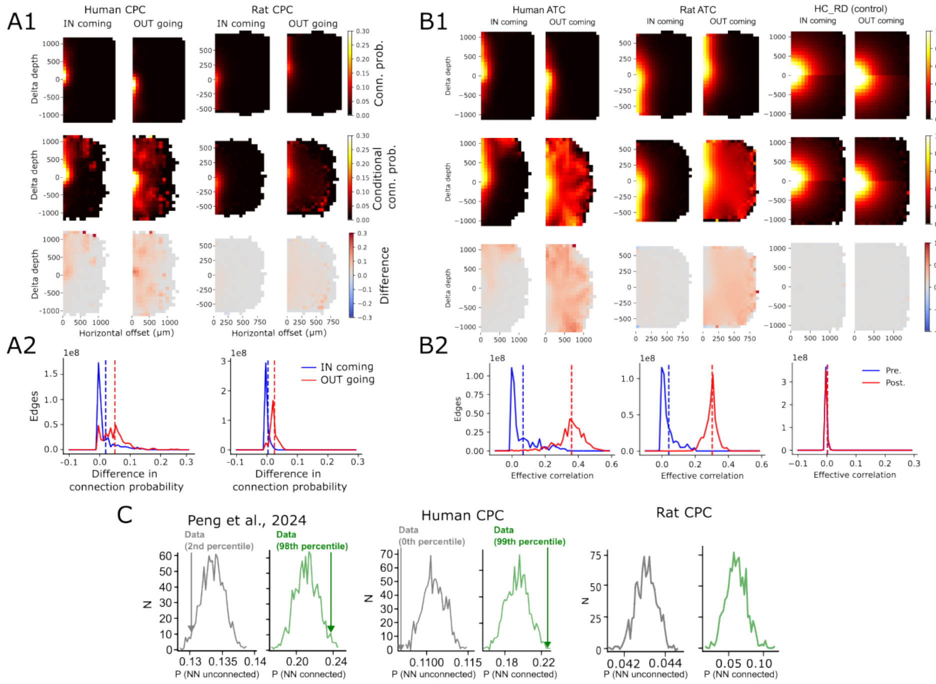


**Figure S3** (A1) Contrast between connection probabilities into a distance bin, with the connection probability conditioned on the nearest neighbor of a neuron being connected. All pairs were considered. Additionally, two-dimensional offset bins are considered instead of distances in one dimension. Overall connection probability in the indicated direction (top),conditioned on the nearest neighbor being connected (middle), difference (bottom). Strength of the effect captured in human (left) and rat (right) CPCs. (A2) Distribution of the difference of connection probability across all edges of the corresponding network, incoming (blue) and outgoing (red) connectivity. Dashed lines indicate respective means. (B1 and B2) Same as A1 and A2, respectively, for ATC networks of human (left), rat (middle) and the HC_RD (right). (C) Distribution of overall and conditional connection probabilities at all distances for 1000 configuration model controls of the indicated networks. The arrows indicate the location of the corresponding value in the actual (non-control) networks.

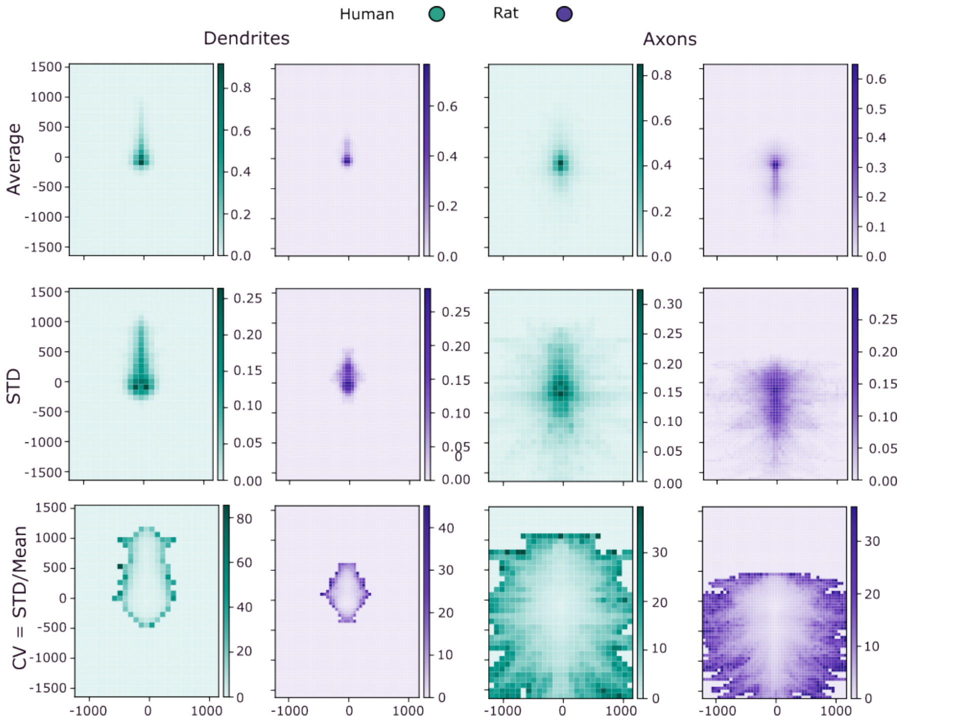


**Figure S4** Density clouds. Dendrites and axons were randomly rotated (20 times) and projected to x-y plane to generate a density cloud of each neuron. From this it was computed: the Average (top), standard deviation (middle) and coefficient of variation (bottom) for dendrites (left) and axons (right)

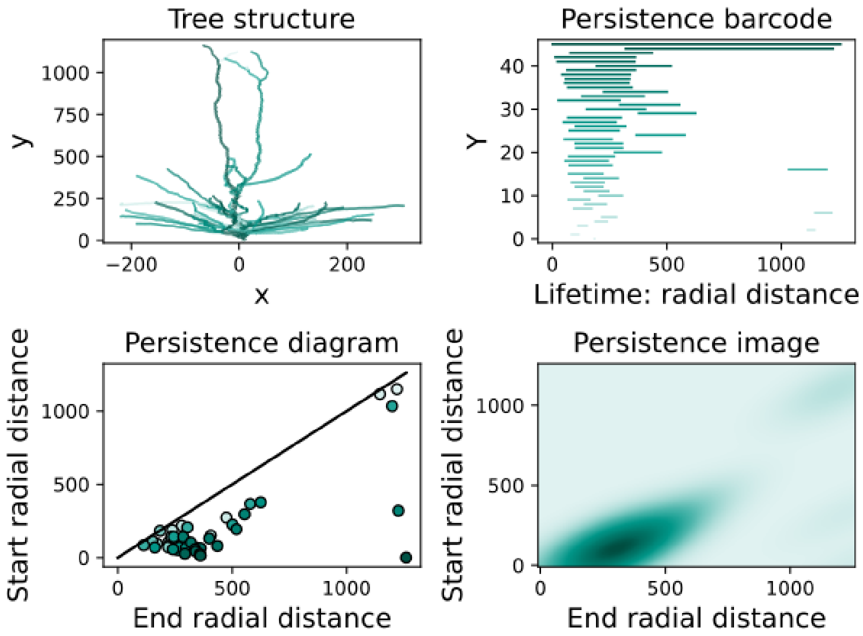


**Figure S5** Dendritic tree structure of an exemplar of a pyramidal neuron (top left). Persistent analysis from **?** ]: bar-code (top right), diagram (bottom left) and image (bottom right).
